# Phospholipid transport to the bacterial outer membrane through an envelope-spanning bridge

**DOI:** 10.1101/2023.10.05.561070

**Authors:** Benjamin F. Cooper, Robert Clark, Anju Kudhail, Gira Bhabha, Damian C. Ekiert, Syma Khalid, Georgia L. Isom

## Abstract

The outer membrane of Gram-negative bacteria provides a formidable barrier, essential for both pathogenesis and antimicrobial resistance. Biogenesis of the outer membrane requires the transport of phospholipids across the cell envelope. Recently, YhdP was implicated as a major protagonist in the transport of phospholipids from the inner membrane to the outer membrane however the molecular mechanism of YhdP mediated transport remains elusive. Here, utilising AlphaFold, we observe YhdP to form an elongated assembly of 60 β strands that curve to form a continuous hydrophobic groove. This architecture is consistent with our negative stain electron microscopy data which reveals YhdP to be approximately 250 Å in length and thus sufficient to span the bacterial cell envelope. Furthermore, molecular dynamics simulations and in vivo bacterial growth assays indicate essential helical regions at the N- and C-termini of YhdP, that may embed into the inner and outer membranes respectively, reinforcing its envelope spanning nature. Our in vivo crosslinking data reveal phosphate-containing substrates captured along the length of the YhdP groove, providing direct evidence that YhdP transports phospholipids. This finding is congruent with our molecular dynamics simulations which demonstrate the propensity for inner membrane lipids to spontaneously enter the groove of YhdP. Collectively, our results support a model in which YhdP bridges the cell envelope, providing a hydrophobic environment for the transport of phospholipids to the outer membrane.

## Introduction

The tripartite cell envelope of Gram-negative bacteria provides a formidable barrier against harmful exogenous substances, such as antibiotics, and thus is a major contributor to intrinsic antimicrobial resistance. The cell envelope consists of an inner membrane (IM) and outer membrane (OM) separated by an aqueous periplasm. The OM is asymmetric in nature, with phospholipids (PLs) and lipopolysaccharide (LPS) populating the inner and outer leaflets respectively, and adorned with an array of proteins (1). To build and maintain this complex OM structure, various hydrophobic constituents must be transported across the aqueous periplasm. Thus far, several proteinaceous transport systems have been demonstrated to orchestrate this process, either by shuttling lipids/proteins between membranes via soluble carrier proteins (2–4), or assembling large periplasmic-spanning systems that directly connect the inner and outer membranes (5, 6).

To date, a major unknown in OM biogenesis is the mechanism by which PLs are transported to the OM. The most extensively characterised intermembrane PL transport pathway remains the *E. coli* Maintenance of Lipid Asymmetry (Mla) pathway (2, 7), which consists of a periplasmic protein that shuttles PLs between an IM ABC transporter and an OM complex (8–19). However, whilst some studies indicate a role in anterograde transport (20, 21), the majority of research suggests the Mla pathway facilitates retrograde transport, removing mislocalised PLs from the outer leaflet of the OM (2, 10, 14, 15, 17, 22, 23).

Indeed another protein, YhdP, has recently emerged as a potential anterograde PL transporter (24, 25). This arose during a screen for genetic interactions of MlaA*, a gain of function mutation in MlaA, an OM component of the Mla pathway. MlaA* causes increased transit of PLs into the outer leaflet of the OM (25, 26), causing destabilisation of the OM and increased OM vesicle production. The *mlaA\** cells compensate for this destabilisation by elevating anterograde PL flux from the IM to the OM. This increased flux can be modulated by loss of function mutations in *yhdP*, suggesting the involvement of YhdP in anterograde PL trafficking (25).

YhdP is one of six paralogous AsmA-like proteins found in the *E. coli* cell envelope, which have been recently reviewed by Kumar and Ruiz, 2023 (27). All six proteins are predicted to be anchored to the IM and extend into the periplasm (28, 29), well situated for transport across the cell envelope. Strains lacking *yhdP* possess phenotypes indicative of a defective OM (24, 30), which are exacerbated in a *yhdP tamB* double knockout strain, resulting in altered cell morphology and significant sensitivity to antimicrobials (30, 31). Moreover, *yhdP* depletion in combination with the removal of *tamB* and *ydbH* induces an accumulation of PLs in the IM, presumably due to defective anterograde PL transport (30). Importantly, the simultaneous deletion of *yhdP, tamB* and *ydbH* results in the loss of cell viability (30, 31), implying their involvement in an essential cell process.

Beyond *E. coli*, AsmA-like proteins are widespread in the cell envelopes of Gram-negative bacteria, and have been linked to virulence in pathogens (32), which, in combination with their role in antimicrobial resistance (25, 30, 31), pose AsmA-like proteins as attractive therapeutic targets. Furthermore, AsmA-like proteins likely share a common ancestor with transport systems widespread across eukaryotes (33, 34), which have emerged as bridges through which lipids from neighbouring organelles may be transported (35–37). Consequently, it is anticipated that bacterial AsmA-like proteins, and their eukaryotic homologues, likely perform a common role in inter-membrane lipid transport.

Collectively, these existing data support the notion that YhdP functions as an anterograde PL transporter, acting to build and maintain the OM. Nevertheless, the mechanism by which YhdP functions at the molecular level remains fundamentally uncertain. Here, we present data to support a model in which YhdP forms an elongated groove across the periplasm that directly connects the inner and outer membranes, through which PLs are transported to build and maintain the bacterial OM.

## Results

### YhdP forms a long thin rod consisting of stacked β strands

The gene *yhdP* encodes one of the largest proteins from the *E. coli* genome, a 1266 aa residue (139 kDa) polypeptide. *E. coli* strains lacking *yhdP* exhibit distinct phenotypes indicative of OM defects, such as sensitivity to vancomycin and SDS+EDTA (Supplementary Figure 1A, (24)); that are alleviated by trans expression of *yhdP* from a complementary plasmid. These data are consistent with YhdP facilitating OM biogenesis (24, 25, 30, 31). YhdP is expected to be anchored to the IM via an N-terminal transmembrane (TM) helix, with the bulk of the protein extending into the periplasm (Figure 1A, Supplementary Figure 1B) (28). Intriguingly, a putative TM region is also predicted at the C-terminus of YhdP, but with much lower confidence (Supplementary Figure 1B), thus YhdP may conceivably provide a physical link between both bacterial membranes. YhdP also contains two periplasmic domains (DUF3971 and Asma_C) (Figure 1A), both of which are members of the AsmA-like Pfam domain clan (CL0401) (38). These domains represent regions of sequence homology that are collectively conserved amongst all bacterial AsmA-like family members (38). Furthermore, the N-terminus of YhdP contains a region of sequence homology to the Chorein-N domain present within eukaryotic lipid transfer proteins (33).

**Figure 1.**
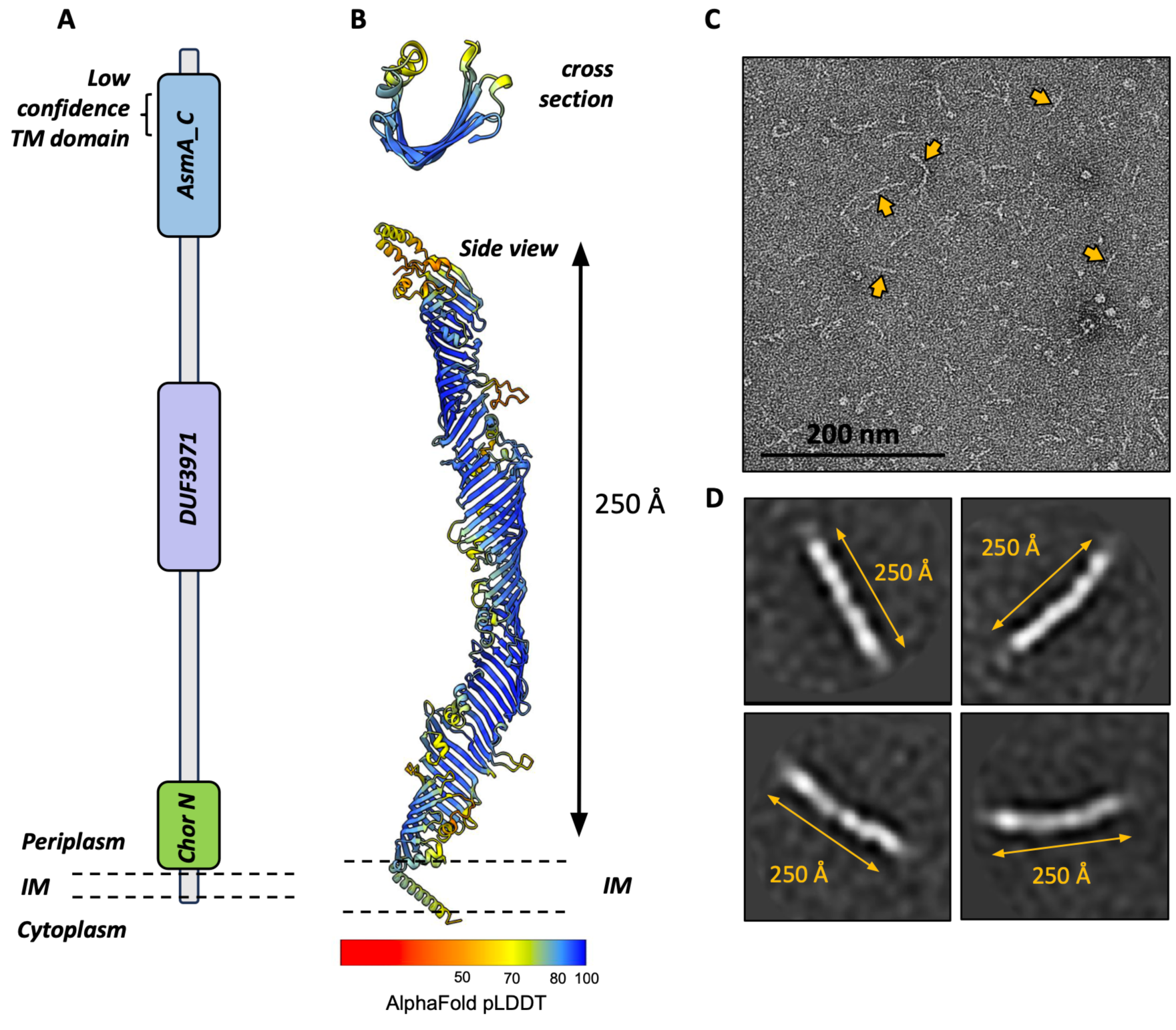
YhdP forms an elongated, thin structure predicted to consist of stacked β strands. (A) Schematic of YhdP, which is predicted to anchor to the inner membrane via an N-terminal transmembrane helix. The region coloured in green reflects that homologous to the eukaryotic Chorein-N domain. The purple and blue regions represent AsmA-like protein domains found within YhdP, DUF3971 and AsmA_C, respectively. At the C-terminus of YhdP a transmembrane helix is predicted at low confidence. (B) AlphaFold 2 model for YhdP, in both a side view and top view/cross section representation, with pLDDT model confidence coloured from red to blue. (C) A negative stain electron microscopy micrograph of purified YhdP. Example YhdP molecules are highlighted with yellow arrowheads. (D) Representative 2D class averages of YhdP negative stain data.

To better understand how YhdP may function as a transport protein, we investigated its architecture. We utilised the AlphaFold structural prediction of YhdP to explore its expected secondary structural elements (29). The per-residue confidence scores associated with the prediction were high for the majority of the model (87 % and 47 % of residues with pLDDT values >70 and >90 respectively) with regions of lower confidence mainly clustered towards the C-terminus (Figure 1B). The AlphaFold model predicts that YhdP adopts an elongated architecture, ∼250 Å in length.

To assess the overall shape of YhdP experimentally, we isolated purified YhdP protein for negative stain electron microscopy. Recombinantly expressed YhdP was isolated from the *E. coli* membrane fraction, demonstrating that YhdP is indeed a membrane protein, yielding a polypeptide of the expected 139 kDa (Supplementary Figure 1C). We imaged YhdP by negative stain electron microscopy, and observed that the protein assembles long, thin rod-type structures (Figure 1C). Subsequent 2D class averages allowed us to estimate the length of YhdP to be approximately 250 Å (Figure 1D), reinforcing our confidence in the validity of the AlphaFold model.

The YhdP AlphaFold prediction constitutes a backbone of 60 stacked β-strands that curve inwards to form a groove (Figure 1B), consistent with the β-taco fold first described in the X-ray structure of a TamB fragment, a paralog of YhdP (Supplementary Figure 2, (39)). This groove appears to extend along the length of YhdP, reminiscent of the Lpt bridge for LPS transport (5), whilst the entire protein twists along its axis. The continuity of β-strands is regularly interrupted by short helical or disordered loop regions along the length of YhdP, a pattern described as the repeating β-groove (RBG) domain in eukaryotic AsmA-like homologs (40), though in bacteria their periodicity appears less defined.

The β-strand dominated architecture of YhdP is reflected in the AlphaFold predicted structures for the remaining five *E. coli* AsmA family members, which vary predominantly in length (Supplementary Figure 2). It should be noted that the AsmA-like Pfam domains representing regions of sequence homology between the *E. coli* AsmA-like proteins (i.e. domains “DUF3971” and “AsmA_C” in YhdP; “TamB” domain in TamB, “DctA-YdbH” domain in YdbH; and the “AsmA” domain in AsmA, YhjG, and YicH (38)) do not form structurally distinct regions (Supplementary Figure 2). Indeed, these domains are indistinguishable as distinct modules within the β-sheet construction of these proteins, implying that the homology between AsmA-like proteins extends far beyond the domains currently annotated.

The structural predictions of bacterial AsmA-like proteins also resemble homologs found in distantly related bacteria and eukaryotes (Supplementary Figure 3, (40)). The Chorein-N domain represents a small homologous region detected between the eukaryotic and bacterial AsmA-Like proteins, based on their amino acid sequences. However, their predicted structures are remarkably similar, implying a substantial structural conservation across life. Indeed, the eukaryotic AsmA-like homologues are also predicted to form long, thin, twisting grooves, further indicating the homology between these groups is far more extensive than sequence homology suggests, and not simply limited to the Chorein-N domain (Supplementary Figure 3). Overall, predicted structures for both the bacterial and eukaryotic proteins highlight the potential limitation of relying solely upon sequence based homology detection and indicate the β-taco fold to be exceptionally conserved throughout evolution, potentially as a module for inter-membrane transport.

### YhdP has the potential to bridge the inner and outer membranes, spanning the periplasm

Periplasmic-spanning systems are emerging as a common means for bacteria to transport hydrophobic components between the inner and outer membranes (5, 6, 41, 42). Our negative stain EM data shows that YhdP is ∼250 A in length, a dimension reinforced by the AlphaFold prediction. YhdP is therefore of sufficient size to span the 210-240 Å periplasm (43–45), directly linking the inner and outer membranes. This model would require the N- and C-termini of YhdP to interact with the IM and OM, respectively. In addition to the TM helix at the N-terminus, we observed other helical regions towards the N- and C-termini of the YhdP AlphaFold prediction that may conceivably form membrane interactions (Figures 2A+D).

**Figure 2.**
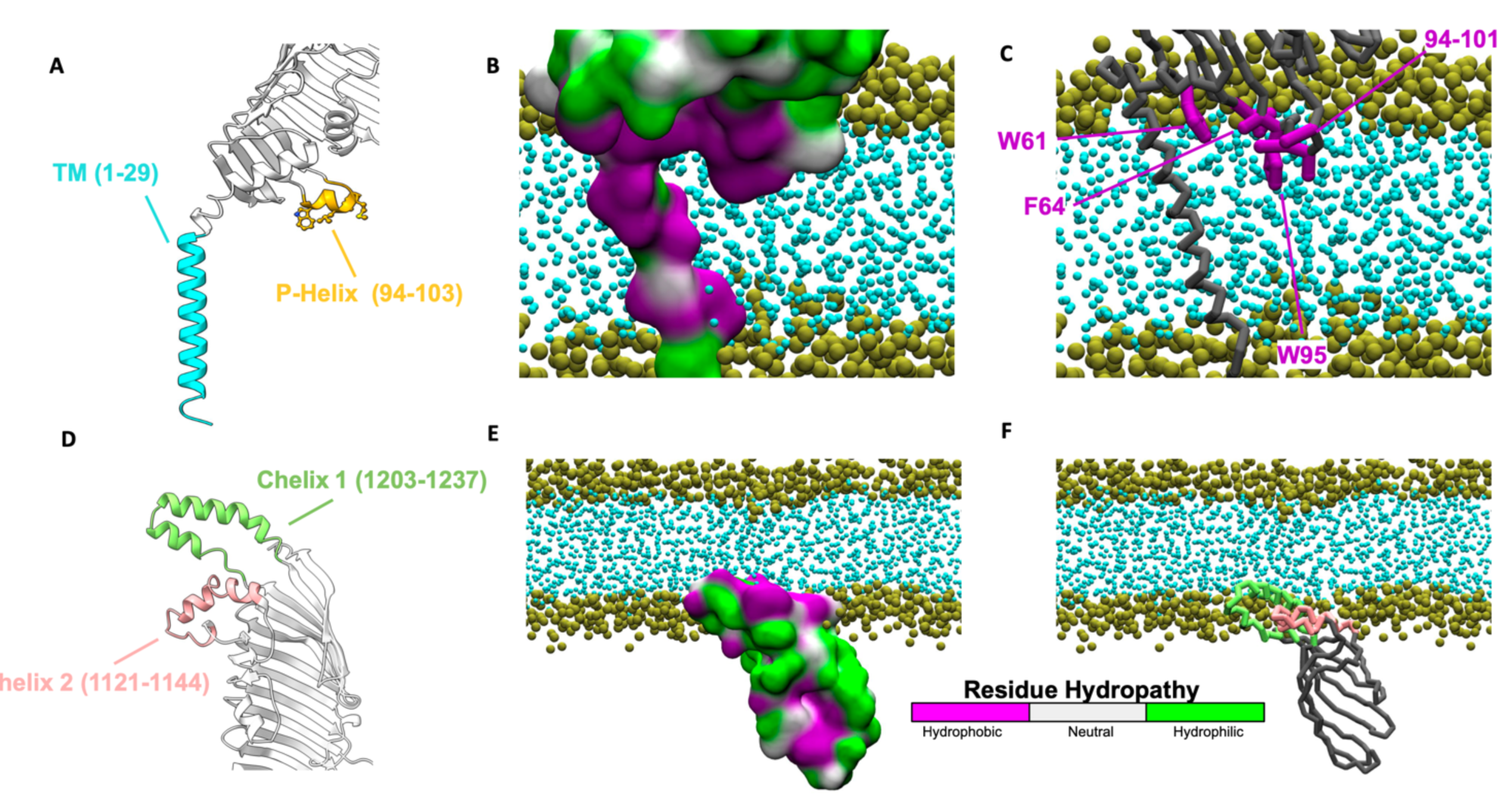
Molecular dynamics simulations suggest that helices at the N- and C-termini of YhdP interact with the inner and outer membranes. (A) Cartoon representation of the N-terminus of the YhdP AlphaFold model. The TM domain and P-helix are highlighted in blue and yellow respectively. (B-C) Representative snapshots from 3 x 5 μs of an N-terminal YhdP fragment (residues 1-387) in a PL bilayer. (B) Surface representation of the N-terminal fragment, coloured by residue hydropathy. (C) Ribbon representation of YhdP with key membrane inserting residues shown as sticks. (D) Cartoon representation of the C-terminus of YhdP. Chelix_1 (residues 1203-1237) and Chelix_2 (residues 1121-1144) are highlighted in green and pink respectively. (E-F) Representative structures from 6 x 5 μs of membrane self-assembly with a C-terminal fragment of YhdP (residues 1098-1266). (E) Surface representation coloured by residue hydropathy revealing the two C-helices form a largely hydrophobic patch. (F) Ribbon representation of the C-terminal fragment revealing that residues of the C-helices remain associated within one leaflet of the membrane. Side chains are coloured thus: strongly hydrophobic residues are purple, residues which are neutral in terms of hydropathy are grey, and hydrophilic residues are green. The backbone is shown in dark grey. Headgroup beads are shown in gold and acyl beads in cyan.

To investigate potential membrane interactions of the N- and C-terminal regions of YhdP, we performed a series of coarse-grained molecular dynamics simulations. First we sought to investigate interactions between the N-terminal region of YhdP and the IM. Here we utilised an N-terminal fragment of YhdP (residues 1-387) which was inserted into a model IM with a representative lipidic composition comprising 1-palmitoyl-2-oleoyl phosphatidylethanolamine (POPE), 1-palmitoyl-2-oleoyl phosphatidylglycerol (POPG) and cardiolipin 2-(CDL2) in a 7:2:1 ratio. We performed three simulations, each of 5 μs duration. Inspection of the trajectories revealed substantial protein interactions with the membrane which, interestingly, were not confined solely to the TM region. The N-terminal entrance of the groove consists largely of hydrophobic residues and sits upon the surface of the membrane (Figure 2B). Many of these hydrophobic residues insert into the membrane, and form interactions with the lipid tails (Figure 2B+C). A small amphipathic helix (residues 94-103) situated between the β3 and β4 strands, henceforth referred as the P (parallel)-helix, sits especially deep within the membrane (Figure 2A-C). The residues on the lower face of the P helix, along with two residues (W61 and F64) within the loop region separating the first two β strands, form a line of hydrophobic residues that appear to stabilise the membrane interaction (Figure 2C). In particular, W95 of the P-helix embeds deeply in the membrane (Figure 2C) and has the highest percentage occupancy (68.6%) of any residue with the lipid tail moieties (Supplementary Figure 4A). These results suggest that the P-helix partially resides within the hydrophobic core of the inner membrane.

Next, we considered interactions between the C-termini of YhdP and the OM, utilising a second truncated fragment comprising the final 169 residues of YhdP (residues 1098-1266). Unlike the N-terminus, where an obvious TM domain is present, it was unclear if or how the C-terminus could interact with a lipid bilayer. Therefore, we employed coarse-grained simulations to investigate membrane self-assembly in the presence of the YhdP C-terminus. Simulations were initiated from randomly placed POPE, water and neutralising ions around the C-terminal YhdP fragment, enabling determination of the preferred protein orientation and localisation with respect to the self-assembled bilayer with minimal bias. We performed six simulations, each of 5 μs duration. In each case a stable lipid bilayer assembled, interacting with the C-terminal end of the hydrophobic groove and two C-terminal helical regions of YhdP (Figure 2E+F). Both C-terminal helices (Chelix_1 - residues 1202-1238, Chelix_2 - residues 1120-1145, Figure 2D) are amphipathic, though the most C-terminal of the two (Chelix_1) is primarily hydrophobic. Indeed, Chelix_1 corresponds with the aforementioned second, putative TM region (predicted with much lower confidence by the TMHMM - 2.0 server (Figure 1A, Supplementary Figure 1B)). In our simulations, Chelix_1 was consistently repositioned to reside closer to Chelix_2, such that the centre of mass separation between the helices was reduced to ∼14 Å, in all but one simulation, having started at a distance >20 Å (Supplementary Figure 5A+B). At the end of the simulations, the hydrophobic groove was positioned perpendicular to the plane of the bilayer (Figure 2E). The C-terminal membrane-interacting regions coalesced to form a large, mostly hydrophobic patch to adhere to the membrane (Figure 2F). Membrane insertion was limited to the inner leaflet of the bilayer, occurring predominantly via the Chelix_1 and Chelix_2 moieties, both of which were inserted into the membrane (Figure 2F). Indeed, Chelix_1 and Chelix_2 accounted for 49.5% and 24.6% of all protein-lipid tail interactions, respectively, with interactions between these helices and the POPE acyl tails dominated by hydrophobic residues (Supplementary Figure 4B).

Finally, we investigated whether the membrane interactions we had identified using N- and C-terminal protein fragments were sufficient to anchor YhdP within the context of the Gram-negative cell envelope. We simulated the entire system, with each TM region embedded in an appropriate OM or IM model membrane, for 3 simulations of 2 μs. We utilised our previously published *E. coli* cell envelope system containing the AcrAB-TolC efflux system to define the spacing between the two membranes (Supplementary Figure 6) (46). YhdP remained stably bound throughout the simulation, displaying membrane interactions at both termini in this full system similar to those previously observed with the smaller protein fragments (Supplementary Figure 4C+D, Supplementary Video 1). We did not observe any interaction between YhdP and the outer leaflet of the OM, strengthening our hypothesis that the C-terminus only interacts with the inner leaflet of the OM.

### Putative membrane-interacting helices at the N and C-termini are essential for YhdP function

Collectively, our negative stain EM data, the AlphaFold model, and our molecular dynamics simulations support a model where YhdP forms an envelope spanning groove, that may directly interact with the inner and outer membranes, via helical regions at its N- and C-termini (Figure 3A). We therefore sought to further investigate the importance of these helical regions by assessing the functionality of YhdP mutants with these helices removed. Consequently, we utilised the vancomycin and SDS+EDTA sensitivity of a *yhdP* knockout, as described above (Supplementary Figure 1, (24)), to evaluate the ability of *yhdP* mutants to alleviate these defects in a phenotypic complementation assay (Figure 3B).

**Figure 3.**
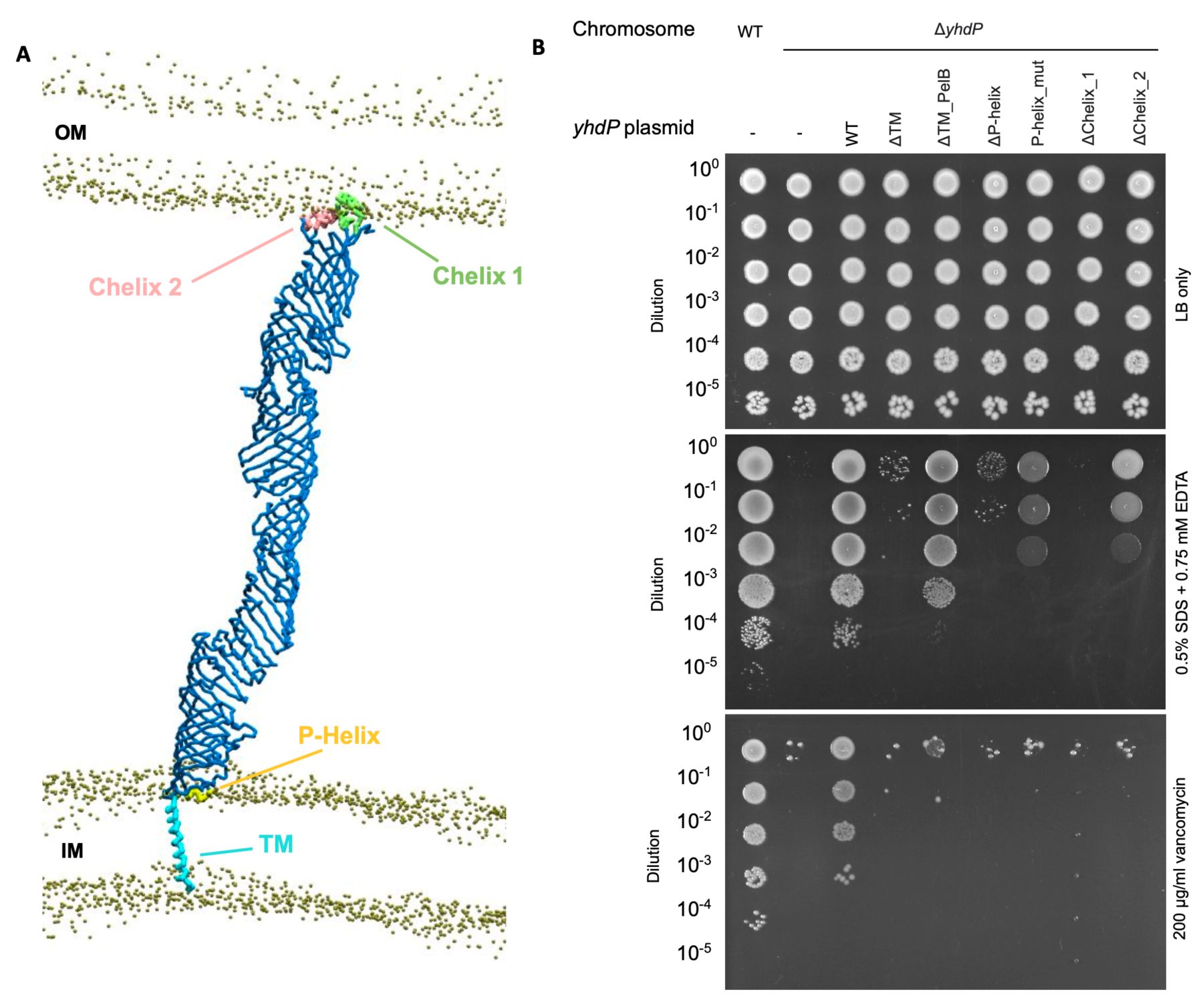
The N- and C-terminal helices of YhdP are required for function. (A) A snapshot of YhdP simulated in the context of two membranes with the key helices at the N- and C-termini highlighted. Backbone beads are shown in blue, helices are coloured as labelled, and phosphate beads are shown in gold. Other lipid beads are omitted for clarity. (B) Genetic complementation of an *E. coli* K-12 *yhdP* knockout strain with different plasmid encoded mutants of YhdP. 10-fold serial dilutions of the indicated cultures spotted on LB plates containing SDS+EDTA or vancomycin at the concentrations indicated.

First, we investigated the requirement of YhdP to interact with the IM at its N-terminus. We targeted the N-terminal TM helix by constructing two mutants; one with the TM domain deleted (ΔTM), resulting in presumed cytoplasmic retention of YhdP, and another where the TM domain was replaced with a PelB leader sequence (ΔTM_PelB), for secretion of YhdP into the periplasm. In the presence of vancomycin, neither of these mutants could complement the *ΔyhdP* phenotype (Figure 3B). In contrast, while the ΔTM mutant was unable to complement in the presence of 0.5% SDS+0.75 mM EDTA, the ΔTM_PelB mutant almost fully restored growth (Figure 3B). Nevertheless, increasing the amount of EDTA further decreased the growth of the ΔTM_PelB mutant relative to the WT, suggesting it is still less functional than WT YhdP (Supplementary Figure 7). Collectively, these data indicate that the periplasmic localisation of YhdP is essential for its function but that some functionality is retained upon removal of its N-terminal TM region.

In addition to the TM domain, we also targeted the N-terminal P-helix (Figure 2A+Figure 3A). To assess the importance of this helix, we constructed two further mutants; one where the entire P-helix was deleted (ΔP-helix), and a second with the four hydrophobic helix residues mutated to polar or charged amino acids (P-helix_mut). Neither mutant could restore growth of the *ΔyhdP* strain, though the P-helix_mutate mutant grew slightly better than ΔP-helix on SDS+EDTA (Figure 3B). Surprisingly, these results suggest that the P-helix has a larger impact upon YhdP function than its TM helix. Indeed, its embedding within the IM may explain the ability of YhdP to remain partially functional without its TM helix.

Considering this possibility, we utilised molecular dynamics to assess YhdP-IM interactions in the absence of the TM region. We performed 3 x 5 µs coarse-grained simulations starting with YhdP in the same position as the original N-terminal simulations, but with the TM domain (residues 2-31) removed. The resulting Δ2-31 N-terminal fragment remained associated with the membrane throughout our simulations (Supplementary Figure 8A). Indeed, this observation corroborates our hypothesis that the P-helix alone is sufficient to maintain YhdP’s IM interaction and reinforces its importance for YhdP functionality. As previously, we analysed the protein-lipid contacts and observed a similar pattern of lipid tail interactions, with W95 once again displaying the highest lipid acyl tail occupancy (67.7%) (Supplementary Figure 8B+C). Overall, we anticipate the P-helix to be particularly important in stabilising the YhdP-IM interaction; however, considering YhdP’s possession of a canonical TM region, it seems unlikely that the P-helix exists purely to mediate IM interactions. Indeed, this notion is reinforced by the substantial decrease in the ability of YhdP to complement the *ΔyhdP* phenotype upon removal or mutation of the P-helix in constructs retaining their TM regions.

Finally, we probed the C-terminal helices of YhdP, regions that may interact with the OM and/or other proteins within the OM. We observed that deletion of either ΔChelix_1 or ΔChelix_2 prevented complementation of the *ΔyhdP* phenotype on vancomycin, though ΔChelix_2 grew better than ΔChelix_1 in the presence of SDS+EDTA (Figure 3B). These results indicate that both C-terminal helical regions of YhdP are required for function, with Chelix_1 appearing essential, potentially due to its predicted ability to embed in the inner leaflet of the OM (Figure 2F). To support these observations, we performed three molecular dynamics simulations of randomised bilayer assembly in the presence of a YhdP C-terminal mutant lacking Chelix_1 (residues 1098-1266, replacing residues between E1202 and S1238 with GS repeats). The three simulations each resulted in rather different protein-membrane binding modes, none of which permitted a productive interaction between the YhdP groove and the membrane, suggesting Chelix_1 is required for stable attachment of YhdP to the OM during our simulations (Supplementary Figure 8D). These molecular dynamics simulations, together with our phenotypic complementation assays, support a model whereby the C-terminal helices of YhdP, particularly Chelix_1, are required for YhdP function, potentially in anchoring YhdP to the OM.

Indeed, whilst further investigations are required to fully investigate the functions of the helical regions at the N- and C-termini of YhdP, our data thus far supports a model in which YhdP interacts with both membranes, thus spanning the periplasm.

### Phospholipids traverse the groove of YhdP

Given YhdP is anticipated to transport PLs (25, 30, 31), we inspected its predicted structure for insight into how such transport may be mediated. The most pronounced feature of YhdP is a groove, which using the MOLE*online* server (47) was revealed to be continuous along all but the extreme N-terminus of the protein (Figure 4A). Visualisation of the molecular lipophilicity potential and Coulombic electrostatic potential of YhdP revealed the groove interior to be hydrophobic and uncharged, whilst the periplasmic-facing exterior face is hydrophilic and predominantly negatively charged (Figure 4B+C). This predicted groove presents an ideal environment for shielding the acyl tails of PLs, or other hydrophobic moieties, whilst their hydrophilic portions remain exposed to the aqueous periplasm.

**Figure 4.**
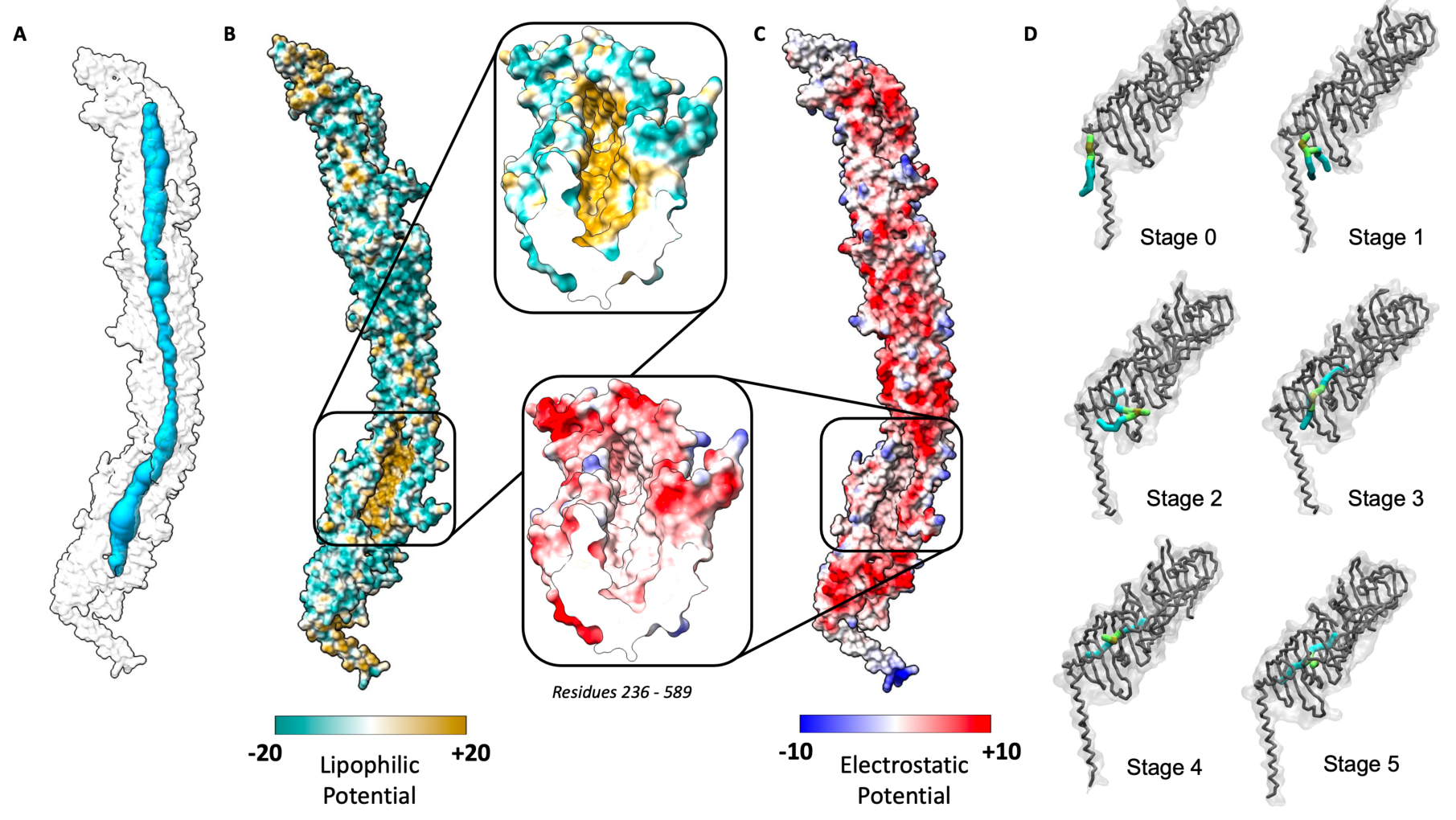
YhdP AlphaFold model contains a continuous hydrophobic groove along the majority of its length. (A) Molecular surface representation of the YhdP AlphaFold prediction (white) indicating the existence of a solvent exposed channel along all but the extreme N-terminus of YhdP (blue). (B) Lipophilic potential mapped to a surface representation of the AlphaFold YhdP prediction. Surfaces are coloured from hydrophilic (dark cyan) to hydrophobic (gold). Lipophilicity potential was calculated using the ChimeraX mlp command. Inset (right) depicts a cross section of the YhdP groove highlighting the lipophilicity of its lining. (C) Coulombic electrostatic potential mapped to a surface representation of the AlphaFold YhdP prediction. Surfaces are coloured from positive charge (blue) to negative charge (red). Electrostatic potential was calculated using the ChimeraX coulombic command. Inset (left) depicts a cross section of the YhdP groove highlighting the absence of charge within its lining. (D) Representative surface and backbone snapshots from one of the 5 µs simulations of the N-terminal fragment (residues 1-387). In Stage 0, PLs are in the membrane without any association with YhdP. In Stage 1, PLs rotate their lipid tails towards the mouth of the YhdP hydrophobic groove while lipid headgroups remain away. In Stage 2, an acyl tail explores further up the hydrophobic groove, while the lipid headgroup remains solvated. In Stage 3, the lipid headgroup moves inside the hydrophobic groove, followed by Stage 4 where the lipid headgroup then re-emerges from the hydrophobic groove into a cavity within the surface of YhdP. In stage 5 the lipid moves further within the groove and the headgroup emerges from a second cavity within the surface of YhdP. Lipid colour key: acyl beads in cyan, glycerol beads in lime, phosphate beads in gold. Stick representation of YhdP cavity residues shown in red. Backbone of YhdP shown in grey.

During 8 of our 9 previously described simulations containing the YhdP N-terminus, we observed a single lipid moving into the hydrophobic groove of the protein (Supplementary Video 2), noting that multiple different lipid species entered across our simulations (Supplementary Figure 9). The mechanism of entry into YhdP was consistent in each of our simulations. First, if not already in close proximity to YhdP, lipids in the outer leaflet of the bilayer moved towards the mouth of the groove of YhdP (Figure 4D, Stages 0 -1). Next, the lipid tails tilted away from the membrane towards the groove of YhdP (Figure 4D, Stage 1). The tails then started to explore deeper in the groove of YhdP, while the headgroups remained in the periplasm (Figure 4D, Stage 2). Complete entry of the lipid into the hydrophobic groove then occurred via the following mechanism; an acyl tail extended higher into the hydrophobic groove, followed by the glycerol backbone moieties and finally the polar headgroup, resulting in the lipid adopting a splayed conformation (Figure 4D, Stage 3). The headgroup subsequently emerged in a small periplasmic-exposed cavity, with the acyl chains remaining deep within the groove of YhdP (Figure 4D, Stage 4). In 3 of our simulations, the lipid then travelled further up the groove and the headgroup re-emerged in a second cavity (Figure 4D, Stage 5). Here, the lipid is at the base of a continuous lipid conduit through which it could travel towards the OM (Figure 4A). While the mechanism described here requires further investigation, our molecular dynamics simulations support a model where PLs can enter the groove at the N-terminus of YhdP, to gain access to the lipid conduit.

To directly test if PLs traverse the groove of YhdP, we sought to utilise an *in vivo* crosslinking assay for the capture of phosphate-containing molecules, such as PLs (8, 42). Specifically, we wanted to examine if PLs interact directly with YhdP and, if so, where within YhdP they are located. We incorporated the unnatural, photo-crosslinkable amino acid p-benzoyl-L-phenylalanine (Bpa) into YhdP at 15 different locations. We chose 10 locations along the length of the protein that are predicted to line the aforementioned groove for Bpa substitution: W227, F297, F464, F639, F705, W843, F888, W992, F1070, and Y1085 (Figure 5A). In parallel, we selected 5 residues predicted to face outwards into the periplasm: K264, R663, E873, K960 and K1057.

**Figure 5.**
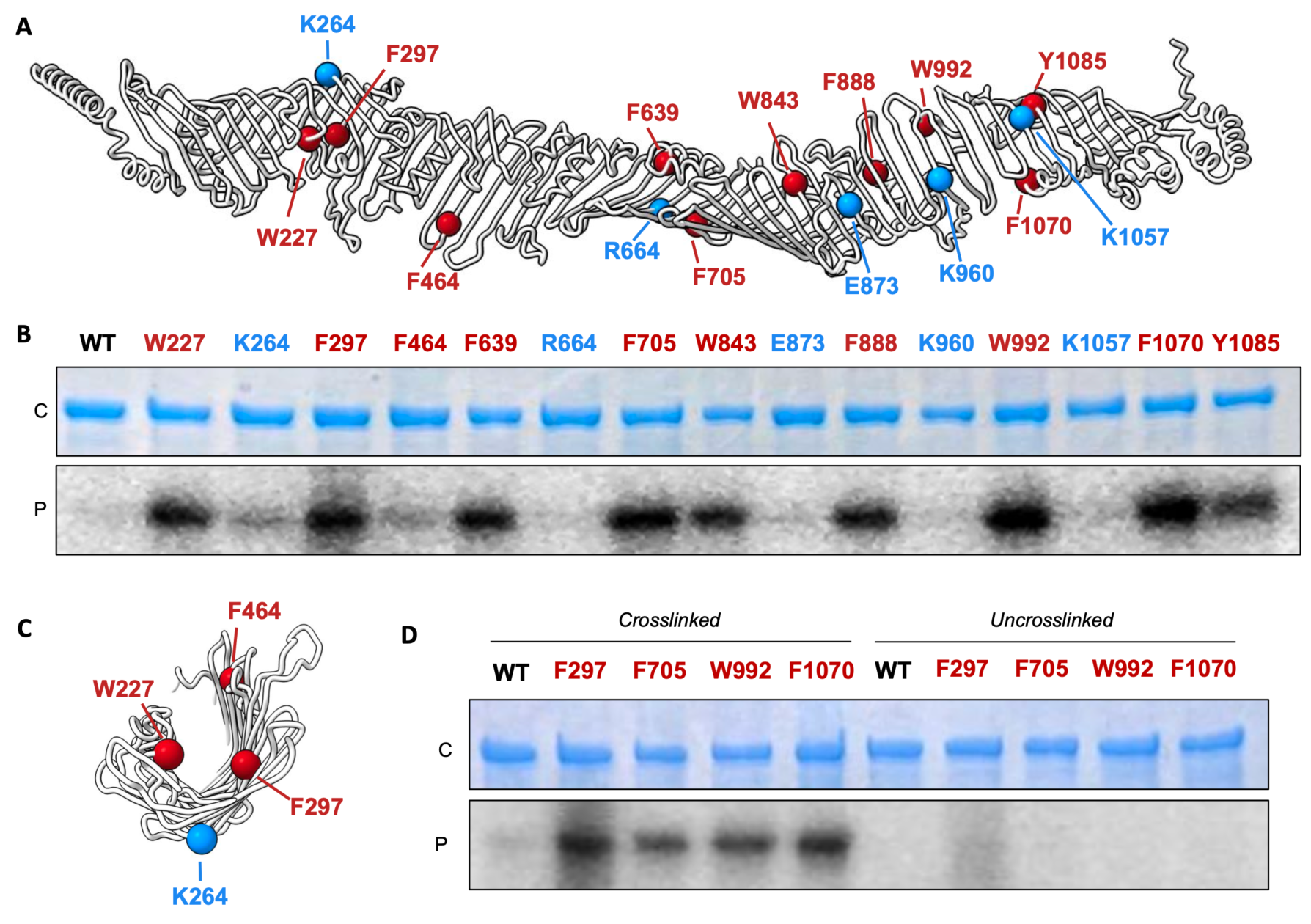
Phosphate-containing molecules in the groove of YhdP. (A) Liquorice cartoon representation of YhdP indicating the sites of Bpa cross-linker incorporation (spheres). Red, residues inside the groove and blue, residues outside the groove. (B) SDS-PAGE of purified and crosslinked WT YhdP and Bpa mutants, visualised by Coomassie staining (protein, C) or phosphorimaging (_32_P signal, P). (C) Cross-section of a segment of YhdP indicating the sites of Bpa cross-linker incorporation (spheres). (D) Uncrosslinked controls of select Bpa mutants inside YhdP. See supplementary figure 10 for uncropped gels.

Wild type YhdP and the 15 mutant proteins were expressed in *E. coli* in the presence of 32P orthophosphate resulting in the bulk labelling of all phosphate-containing molecules, including PLs. After UV treatment, which induces crosslinks between Bpa and nearby molecules, we purified the proteins and analysed them by SDS-PAGE and phosphor imaging. For 9 of the 10 inward facing residues, we observed a very strong phosphor signal (Figure 5B). As Bpa forms crosslinks with C-H bonds, and there is evidence for YhdP as a PL transporter (25, 30, 31), we hypothesise that the modified residues form crosslinks with the acyl chains of PLs as they transit the protein. The single inward-facing residue that formed less crosslinks was F464, which may be explained by its location towards the edge of the groove where it would potentially have less contact with PL acyl chains (Figure 5C).

In contrast to the inward facing residues, the detected phosphor signal was distinctly lower for the outward facing residues, and the WT control, indicating that efficient cross-linking to phosphate-containing substrates required the modified residues to line the groove of YhdP. To ensure the observed signal arose from site specific crosslinking, we selected the 4 inward facing residues yielding the highest phosphor signals (F297, F705, W992, F1070), and performed a separate set of replicates with both crosslinked and uncrosslinked samples (Figure 5D). Whilst the crosslinked samples reflected those previously observed, there was a distinct reduction in the phosphor signal of the uncrosslinked samples confirming that the signals detected reflected Bpa dependent crosslinking to phosphate-containing molecules at these specific locations.

Our Bpa crosslinking assay provides two key insights into the function of YhdP. Firstly, YhdP directly interacts with phosphate-containing molecules, consistent with the genetic data that YhdP functions as a PL transporter (25, 30, 31). Secondly, these data indicate that phosphate-containing molecules, such as PLs, are present along the entire length of the YhdP groove, supporting our hypothesis that PLs would travel along this groove for transport between the inner and outer membranes.

## Discussion

In this study we provide direct evidence to support a model in which YhdP forms a periplasmic-spanning bridge through which PLs are transported to the bacterial OM. We identify that helical regions at the N- and C-terminus of YhdP are functionally important, potentially due to interactions with the inner and outer membranes, respectively. Of interest is the short amphipathic helix (P-helix) that our molecular dynamic simulations indicate to embed within the outer leaflet of the IM. Similar helices have been reported previously in a cryo-EM structure of the IM PL transport complex, MlaFEDB (8), and are implicated in coordinating interactions between protein subunits, suggesting that the P-helix in YhdP could perform a similar role with yet-to-be identified partner proteins. The two helices towards the C-terminus of YhdP we anticipate to directly interact with the inner leaflet of the OM. It is unclear whether such interactions would be purely to anchor YhdP to the OM, or if they somehow aid in OM lipid insertion, either directly or via interaction with OM protein(s). Nevertheless, as OM proteins usually adopt β barrel conformations or are anchored as lipoproteins, the direct interaction of helices with the OM is an unusual observation and one which merits further investigation.

A striking feature of the YhdP Alphafold model is a hydrophobic groove that is continuous along all but the extreme N-terminus of the protein. Our molecular dynamics simulations of YhdP revealed IM PLs to spontaneously enter the groove, in a splayed conformation, with one tail facing upwards towards the OM, whilst the other faces downwards toward the IM. A similar splayed PL conformation has been previously observed in MlaFEDB (8), suggesting this may be an energetically favourable conformation for PLs to be extracted from or inserted into the membrane. While our simulations suggest that YhdP is amenable to PL uptake, further analyses are required to fully assess the validity of the splayed conformation, and how this may alter in the event of bulk PL transport in which AsmA proteins have been implicated (27). Nevertheless, our *in vivo* cross-linking data indicates the presence of putative PLs throughout the length of the groove, supporting a model in which PLs traverse the length of the YhdP groove for direct transport between the inner and outer membranes (Figure 6). Together, our model supports the previous studies proposing YhdP as the principal anterograde PL transporter in *E. coli* (25, 30, 31).

**Figure 6.**
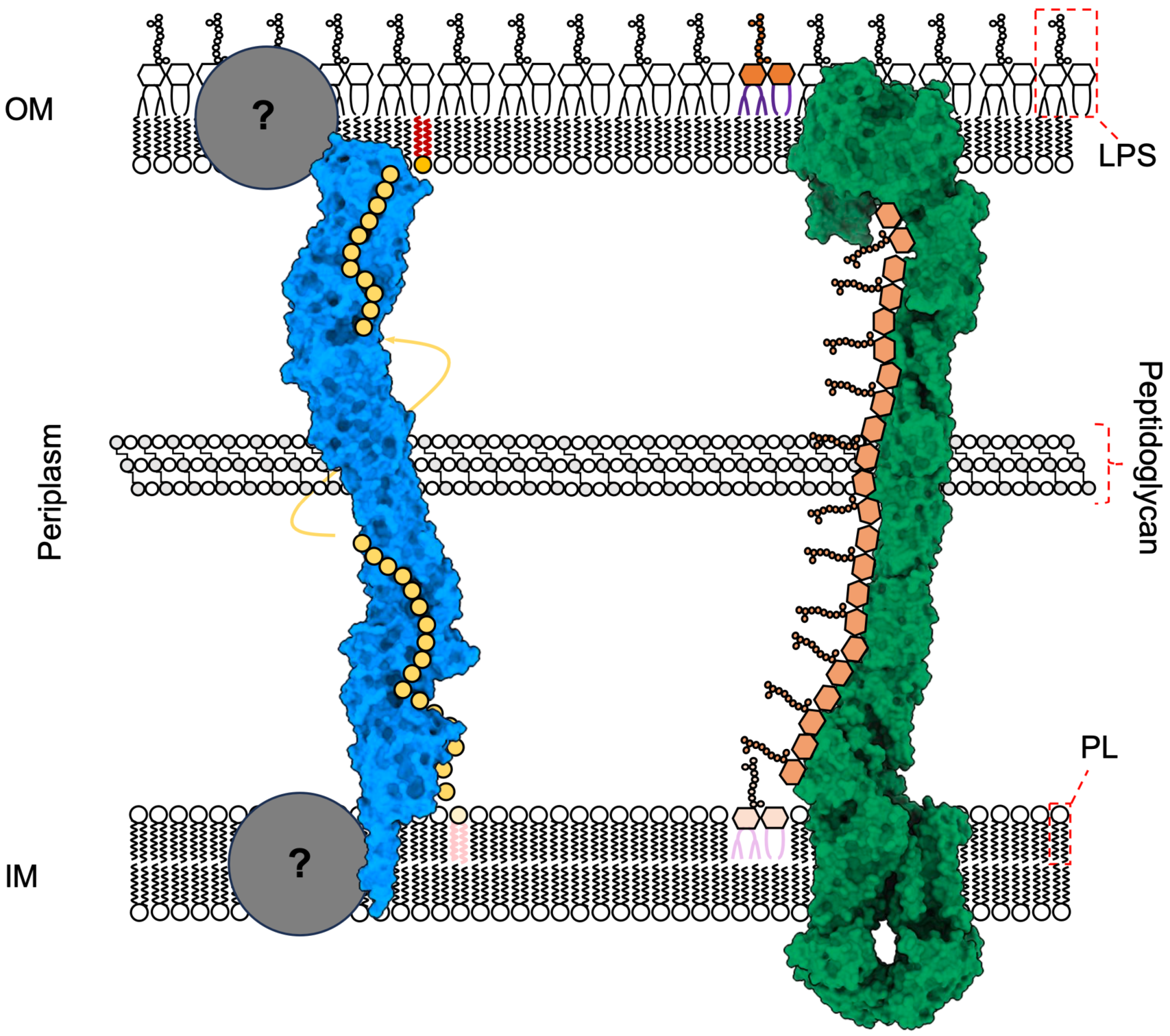
Model for YhdP mediated PL transport in the context of the cell envelope. Surface representation of YhdP (blue) and the Lpt system (green) in the bacterial cell envelope. IM lipids are loaded into the N-terminal of YhdP, potentially assisted by as yet unidentified IM partner proteins. Lipids traverse along the hydrophobic groove of YhdP towards the OM. Once again, the mechanism of unloading at the OM is unclear and is potentially facilitated by additional partner proteins. In contrast, LPS is loaded into the periplasmic bridge structure, formed of LptA monomers, by the IM LptB_2_FGC machinery and sequentially shunted towards the OM. Once at the OM, the insertion of LPS is performed by the LptDE subcomplex. The Lpt system was assembled from existing deposited structures (PBDs: 5IV9, 2R1A, 6MIT).

The observation that YhdP provides a continuous, shielded, lipophilic environment between the inner and outer membranes evokes comparison to the LptA bridge structure of the Lpt system (Figure 6). Transport of LPS to the OM occurs via a periplasmic bridge, constructed of multiple LptA copies (5, 48). Whilst there is no detectable sequence homology between LptA and YhdP, bridge-type transporters appear a logical mechanism for transporting molecules with both hydrophobic and hydrophilic parts through the aqueous periplasm. One important difference between the Lpt system and YhdP is the presence of identified protein binding partners (Figure 6). The IM LptB_2_FGC ABC transporter uses ATP hydrolysis to extract LPS from the IM and transfer it to LptA (49–51). LPS binding sites are always occupied along the length of the LptA periplasmic bridge, meaning LPS insertion at the IM triggers a sequence of movements that displaces and inserts an LPS molecule into the OM, through the OM complex LptDE (5, 48, 52). With the exception of TamB that is known to interact with the OM β barrel, TamA (53), no binding partners are currently reported for YhdP or the remaining *E. coli* AsmA-like proteins. It is plausible that, like LptA, YhdP would require IM binding partner(s) for the extraction of lipids from the IM, perhaps utilising ATP hydrolysis or the proton motive force. However, it is also conceivable that PL transport occurs independently of cellular energy sources and instead responds to the concentration gradient of PLs present across the cell envelope. This notion is supported by the observation that PL transport to the OM, and the YhdP-dependent flow of PLs, can occur in the absence of ATP (25, 54). Furthermore, it is yet to be determined whether YhdP would require an OM interactor for the insertion of PLs into the inner leaflet of the OM.

Of note, the DedA family of proteins are emerging as potential IM interactors of the AsmA-like family based upon the observation that plant homologs of the DedA and AsmA-like proteins directly interact (55–57). The DedA family of proteins are conserved across life, and are implicated in the transport of lipid molecules across membranes. Furthermore, a recent publication implicated a *Bacillus* DedA protein in the flipping of phosphatidylethanolamine across the inner membrane (55). Whilst further analyses are required, it is appealing to propose that proteins distributing PLs across the IM would work in concert with those transporting PLs between the two bacterial membranes.

That YhdP functions as an inter-membrane transport module is a growing conjecture amongst the field. Our results provide a key step in the molecular characterisation of YhdP,

and support a model in which YhdP spans the periplasm providing a shielded hydrophobic environment for the transport of PLs to aid in the building of the bacterial outer membrane.

## Materials and Methods

### Phenotypic assays for *yhdP* mutants in *E. coli*

Knockouts of *yhdP* were constructed in *E. coli* BW25113 by P1 transduction from the Keio collection (58), followed by excision of the antibiotic resistance cassettes using pCP20 (58, 59). Serial dilutions of the strains in 96-well plates were manually spotted (2 µL each) on plates containing LB agar or LB agar supplemented with 0.5% SDS and 0.75–0.95 mM EDTA, or 200 µg/ml Vancomycin and incubated for 16 hr at 37°C. For complementation and/or testing the functionality of the various YhdP mutants, a pET17b-derived plasmids harbouring either WT or mutant forms of *yhdP* (Supplementary Table 1) were transformed into the *yhdP* knockout strain. We found that leaky expression from the T7 promoter was sufficient for complementation of the phenotypes.

### Expression and purification of YhdP

To obtain purified YhdP, pGLI013, which contains *yhdP* with an N-terminal His-tag, was transformed into T7 express *E. coli* (NEB). For expression, overnight cultures T7 express/pGLI013 were diluted 1:50 in LB (Difco) supplemented with carbenicillin (100 µg/mL) and grown at 37°C with shaking to an OD600 of 0.9, then induced by addition of IPTG to a final concentration of 1 mM, and continued incubation overnight at 15°C. Cultures were harvested by centrifugation, and the pellets were resuspended in lysis buffer (50 mM Tris pH 8.0, 300 mM NaCl, 10% glycerol). Cells were lysed by two passes through an Emulsiflex-C3 cell disruptor (Avestin), then centrifuged at 15,000 xg for 30 min to pellet cell debris. The clarified lysates were ultracentrifuged at 37,000 rpm (F37L Fixed-Angle Rotor, Thermo-Fisher) for 45 min to isolate membranes. The membranes were resuspended in membrane solubilization buffer (50 mM Tris pH 8.0, 300 mM NaCl, 10% glycerol, 25 mM DDM) and incubated for 1 hr, rocking at 4°C. The solubilized membranes were then ultracentrifuged again at 37,000 rpm for 45 min, to pellet any insoluble material. The supernatant was incubated with NiNTA resin (GE Healthcare #17531802) at 4°C, which was subsequently washed with Ni Wash Buffer (50 mM Tris pH 8.0, 300 mM NaCl, 10 mM imidazole, 10% glycerol, 0.5 mM DDM) and bound proteins eluted with Ni Elution Buffer (50 mM Tris pH 8.0, 300 mM NaCl, 250 mM imidazole, 10% glycerol, 0.5 mM DDM).

### Negative stain electron microscopy of YhdP

To prepare grids for negative stain EM, the sample was applied to a freshly glow discharged carbon coated 400 mesh copper grids and blotted off. Immediately after blotting, a 2% uranyl formate solution was applied for staining and blotted off. The stain was applied five times for each sample. Samples were allowed to air dry before imaging. Data were collected on a Talos L120C TEM (FEI) equipped with a 4K x 4K OneView camera (Gatan) at a nominal magnification of 73,000x corresponding to a pixel size of 2.00 Å /pixel on the specimen, and a defocus range of 1 – 2 uM underfocus. Data processing was carried out without CTF correction in Relion 3.0 (Fernandez-Leiro and Scheres, 2017). Micrographs were imported, particles were picked manually followed by automated template-based picking. Several rounds of 2D classification were carried out using default parameters.

### Crosslinking phosphate-containing molecules in YhdP

This method was adapted from (8, 42) Isom et al., 2020, and Coudray et cl,. T7express *E. coli* (NEB) were co-transformed with (1) plasmids to express YhdP (either the WT proteins using pGLI014 (C-terminal histidine tagged YhdP), or derivatives of this plasmid expressing Amber mutant YhdP variants for Bpa incorporation; and (2) pEVOL-pBpF (Addgene #31190), which encodes a tRNA synthetase/tRNA pair for the in vivo incorporation p-benzoyl-l-phenylalanine (Bpa) in *E. coli* proteins at the amber stop codon, TAG (60). Bacterial colonies were inoculated in LB broth supplemented with carbenicillin (100 µg/mL), chloramphenicol (38 µg/mL) and 1% glucose, and grown overnight at 37°C. The following day, bacteria were pelleted and resuspended in 32P labelling medium (a low phosphate minimal media: 1 mM Na_2_HPO_4_, 1 mM KH_2_PO_4_, 50 mM NH_4_Cl, 5 mM Na_2_SO_4_, 2 mM MgSO_4_, 20 mM Na_2_-Succinate, 0.2x trace metals and 0.2% glucose) supplemented with carbenicillin (100 µg/mL) and chloramphenicol (38 µg/mL) and inoculated 1:33 in the 10 mL of the same medium. Bacteria were grown to OD 1.0 and a final concentration of 0.2% L-arabinose, 1 mM IPTG, and 0.5 mM Bpa (Bachem, #F-2800.0005), alongside 375 µCi 32P orthophosphoric acid (PerkinElmer, #NEX053010MC) were added and left to induce overnight. The following day, the cultures were spun down and resuspended in 1 mL of PBS, and the cross-linked samples underwent cross-linking by treatment with 365 nM UV in a Spectrolinker for 30 min. Both the cross-linked and uncross-linked cells were then spun down and resuspended in 1 mL of lysozyme-EDTA resuspension buffer (50 mM Tris pH 8.0, 300 mM NaCl, 10 mM imidazole, 1 mg/mL lysozyme, 0.5 mM EDTA, 25 U/mL benzonase) and were incubated for 1 hr on ice. The cells then underwent eight cycles of freeze-thaw lysis by alternating between liquid nitrogen and a 37°C heat block. The lysate was pelleted at 20,000 xg for 15 min, and the pellets were resuspended in 133 µL of membrane resuspension buffer (50 mM Tris pH 8.0, 300 mM NaCl, 10% glycerol and 25 mM DDM), and incubated, shaking, for 1 hr. The sample volume was then increased to 1 mL using 10 mM wash buffer (50 mM Tris pH 8.0, 300 mM NaCl, 10 mM imidazole) and insoluble material was pelleted at 20,000 xg for 15 min. Each supernatant was then mixed with 25 µL of nickel beads (Cytiva, Ni Sepharose 6 Fast Flow) for 30 min. The beads were pelleted at 500 xg for 1 min and the supernatant collected. The beads were then washed four times with 40 mM wash buffer (50 mM Tris pH 8.0, 300 mM NaCl, 40 mM imidazole, 10% glycerol, 0.5 mM DDM) and finally resuspended in 50 µL of elution buffer (50 mM Tris pH 8.0, 300 mM NaCl, 300 mM imidazole, 10% glycerol, 0.5 mM DDM). The samples were then mixed with 5x SDS-PAGE loading buffer, and the beads spun down at 12,000 xg for 2 min. Eluted protein was analysed by SDS-PAGE and stained using InstantBlue Protein Stain (Expedeon, #isb1l). Relative loading of YhdP on the gel was estimated integrating the density of the corresponding bands in the InstantBlue-stained gel in ImageJ (61), and this was used to normalise the amount of YhdP loaded on a second gel, to enable more accurate comparisons between samples. The normalised gel was stained with InstantBlue and 32P signal was detected using a phosphor screen and scanned on a Typhoon scanner (Amersham). At least three replicates of the experiment were performed, starting with protein expression.

### Structure prediction, analysis and interpretation

Full length structures of the bacterial AsmA-like proteins and their eukaryotic homologues were downloaded from the AlphaFold protein structure database (https://alphafold.ebi.ac.uk/) (29, 62) using the following UniProt accession codes: YhdP - P46474, TamB - P39321, YdbH - P52645, YicH - P31433, AsmA - P28249, YhjG - P37645, Mdm31 - P38880 and Atg2 - P53855, Rv0817c - I6WZH9, SHIP164 - A0JNW5. The crystal structures of TamB and Vps13 fragments were downloaded from the PDB using accession codes 5VTG and 6CBC respectively whilst the Vps13 cryo-EM map was downloaded from the EMDB using accession code EMD-21113.

All structures and cryo-EM maps were visualised using UCSF ChimeraX (v1.5) (63, 64). Molecular lipophilic potential was calculated using the *mlp* command with a specified range of -20 to 20. Coulombic electrostatic potential was calculated using the *coulombic* command with a specified range of -10 to 10. The central channel of YhdP was investigated using the MOLE*online* server (https://mole.upol.cz/) in “pore mode”, with a probe radius and interior threshold of 45 and 0.8 respectively, and the resulting json file interpreted in ChimeraX. Homologous regions between the bacterial AsmA-like proteins and other transport proteins were identified using HHpred at the MPI Bioinformatics Toolkit (https://toolkit.tuebingen.mpg.de/) (65). Full length sequences for all proteins of interest were submitted with the default MSA parameters and searched against the PDB (17th April 2023), Pfam (v35), NCBI Conserved domains (v3.19) and SMART (v6.0) databases.

### Molecular dynamics

All coarse-grained simulations were performed using Martini2.2 forcefield and standard Martini water (66) and all atomistic simulations were performed using the CHARMM36m forcefield (67) with TIP3P water (68). All simulations carried out were coarse grained simulations apart from a simulation to construct a model of the mutated Chelix_1. All simulations were performed using the GROMACS simulation suite (versions 2022.4 & 2021.4) (69).

#### System setup

All simulations were performed within a charge neutralised system with the addition of ions corresponding to a solution with a 0.15 M NaCl concentration.

The N-terminal fragment was built by taking the first 387 residues of the YhdP Alphafold2 prediction. The N-terminal fragment was first positioned in a membrane using the OPM server (70), and then coarse-grained and embedded into a membrane using the Martini Maker module (71, 72) from the CHARMM-GUI server (73, 74). The inner membrane model was composed of POPE:POPG:CDL2 at 7:2:1 ratio.The N-terminal fragment without a TM domain was built by adding a single methionine residue to the N-terminal of a fragment composed only of residues 32-387.

The C-terminal fragment was composed of YhdP residues 1098 to 1266. This fragment was coarse-grained and placed in a box of 124x124x124 Å, containing water and counterions. Some of the waters were then randomly replaced with 420 POPE lipids.

The mutated Chelix_1 was modified by replacing all residues between 1203 and 1237 with GS repeats and a starting structure of the mutated Chelix_1 was created using ColabFold (75). The mutated Chelix_1 was then manually combined with the original C-terminal fragment. An all-atom system of the mutated C-terminal fragment was assembled using CHARMM-GUI’s solution builder module. This atomistic system was energy minimised and equilibrated using the default CHARMM-GUI parameters and then simulated within the NPT ensemble for 100 ns. Positional restraints were applied to backbone and sidechain atoms at 10,000 kJ mol^−1^ nm^−2^, excluding Chelix_1. Following production simulation, cluster analysis was used to determine the most common conformation of the mutated Chelix_1. This structure was manually combined with the original C-terminal fragment. This new mutated fragment provided the starting point for assembling the mutated C-terminal coarse-grained system, which followed the same process described previously.

The double membrane system was built by removing all proteins other than AcrAB-TolC from our previously published model (46). This system was energy minimised and equilibrated to allow repacking of the lipids in response to removal of proteins. The box was then replicated in the x dimension yielding a system containing two x AcrAB-TolC complexes. One of these was removed and YhdP was manually placed in the resulting cavity. A nanopore was placed within one of the lipid bilayers to enable exchange of water between the two compartments.

#### Simulations

The velocity rescale (V-rescale) thermostat (77) with a coupling constant of t = 1 was used for all simulations to maintain the system temperature at 323 K. The pressure was maintained at 1 bar using the Parrinello-Rahman barostat (78) with a time constant of t = 12 ps. An elastic network was added to all models using Martinize.py (66) with a dual cut-off of 0.5 and 0.9 nm, and a force constant of 500 kJ mol^−1^ nm^−2^. The exception was AcrAB-TolC of the full system simulation, which used the ElNeDyn Martini forcefield (79), according to the details provided in a previous report (46). All simulations using Martini2.2 used the recommended parameters for treatment of electrostatics (80). Atomistic simulations used the simulation parameters generated by CHARMM-GUI (74).

The full N-terminal fragment was first subjected to the default CHARMM-GUI energy minimisation protocol. Equilibration followed a modified CHARMM-GUI protocol, where the system was initially simulated within the NVT ensemble, followed by simulation within the NPT ensemble, which involved successively increasing the time step from 2 fs to 20 fs, initially using a Berendsen barostat (81) (5.0 ps coupling constant) and then switching to the Parrinello-Rahman barostat. Restraints were lowered successively, starting at 1000 kJ mol^−1^ nm^−2^ on the backbone beads and 200 kJ mol^−1^ nm^−2^ on the lipid headgroups, ending with zero restraints by the fourth equilibration (Supplementary Table 2). Three separate simulations each of 5 µs duration were carried out within the NPT ensemble, each with randomly assigned velocities. The N-terminal fragment without a TM domain was simulated using the same protocol for three separate simulations of 5 µs.

The C-terminal fragment was first subjected to energy minimisation using the steepest descents algorithm. The minimised system was simulated within the NPT ensemble using the V-rescale thermostat and the Parrinello-Rahman barostat as described above. Six separate simulations of 5 µs each were carried out, each with randomly assigned velocities. The mutated Chelix_1 fragment was simulated using the same protocol three separate times for 5 µs each.

The full system used the default CHARMM-GUI energy minimisation protocol albeit for an extended number of steps. The system was then equilibrated within the NPT ensemble as according to Supplementary Table 3, with integration time step successively increasing from 5 fs to 15 fs. Simulation within the NVT ensemble was not used to allow the membrane to fill in holes left by removal of one AcrAB-TolC. Three simulations of 2 µs were carried out, each with randomly assigned velocities.

#### Analysis

Analysis was carried out using PyLipID (82), MDAnalysis (83, 84) using in house scripts, and in-built GROMACS functions. Visualisation and manipulation were carried out using VMD (85) and PyMOL (The PyMOL Molecular Graphics System, Version 2.5 Schrödinger, LLC.; www.pymol.org).

## Acknowledgments

We thank David Sauer and Clare De’Ath, University of Oxford, for providing critical feedback on the manuscript. We would also like to thank for following funding sources: MR/W016672/1 (MRC career development award to G.L.I), 20POST35210202 (American Heart Association, to G.L.I.), R35GM128777 (NIH/NIGMS, to D.C.E.), G.B. is a Pew Scholar in the Biomedical Sciences, supported by The Pew Charitable Trusts (PEW-00033055), S.K. is funded by the EPSRC grant number EP/V030779 and EP/X035603, and R.C. is funded by the Medical Research Council [grant numbers MR/N013468/1, MR/W006731/1]; Magdalen College; and the Department of Biochemistry, Oxford. We also thank NYU Langone’s Microscopy Laboratory (RRID: SCR_017934) and their Cancer Center Support Grant P30CA016087 at the Laura and Isaac Perlmutter Cancer Center. This work was supported in part through the NYU IT High Performance Computing resources, services, and staff expertise. We acknowledge access to High Performance funding via HECBioSim through EP/X035603.

