## Supplementary figures and video legends for "Phospholipid transport to the bacterial outer membrane through an envelope-spanning bridge"

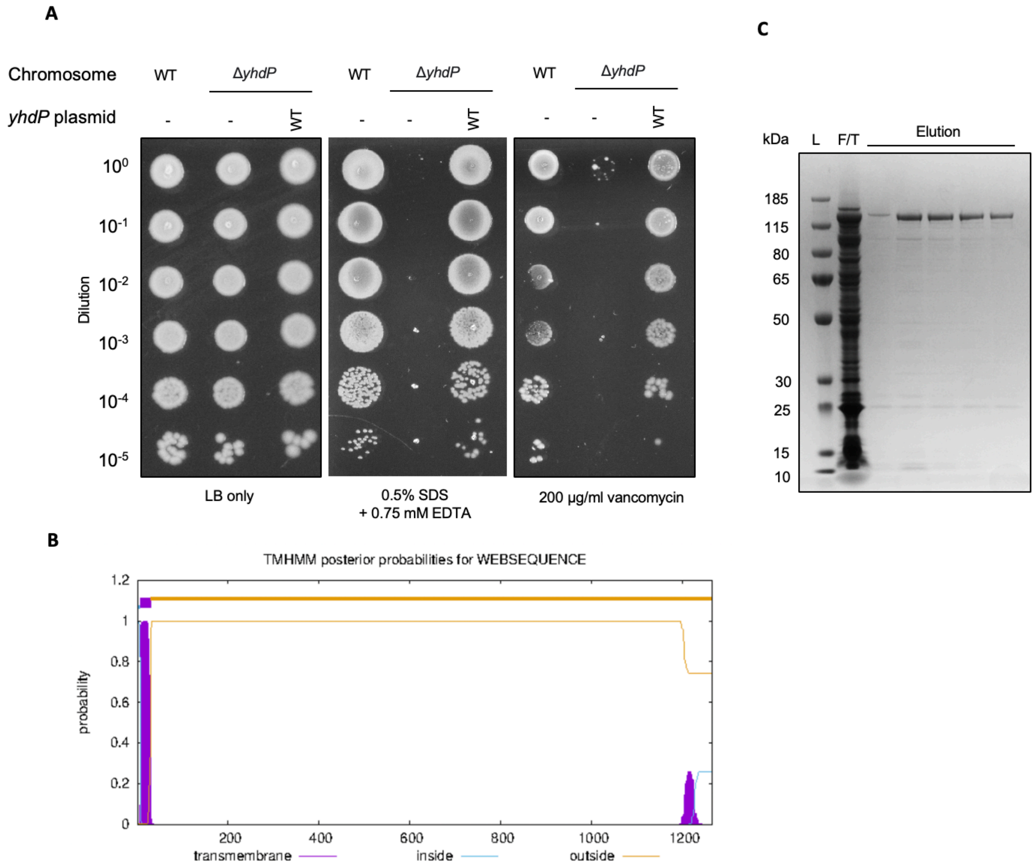


**Supplementary Figure 1 - Phenotypes and purification of YhdP.** (A) Phenotypes of the *yhdP* knockout *E. coli* K-12 strain. 10-fold serial dilutions of the indicated cultures spotted on LB plates containing SDS+EDTA or vancomycin at the concentrations indicated and incubated overnight. The *yhdP* knockout strain does not grow in the presence of SDS+EDTA/vancomycin but can be rescued by the expression of WT YhdP from a plasmid. (B) Prediction of transmembrane domains in YhdP, using TMHMM 2.0. (C) SDS-PAGE of recombinantly expressed and purified his-tagged YhdP.

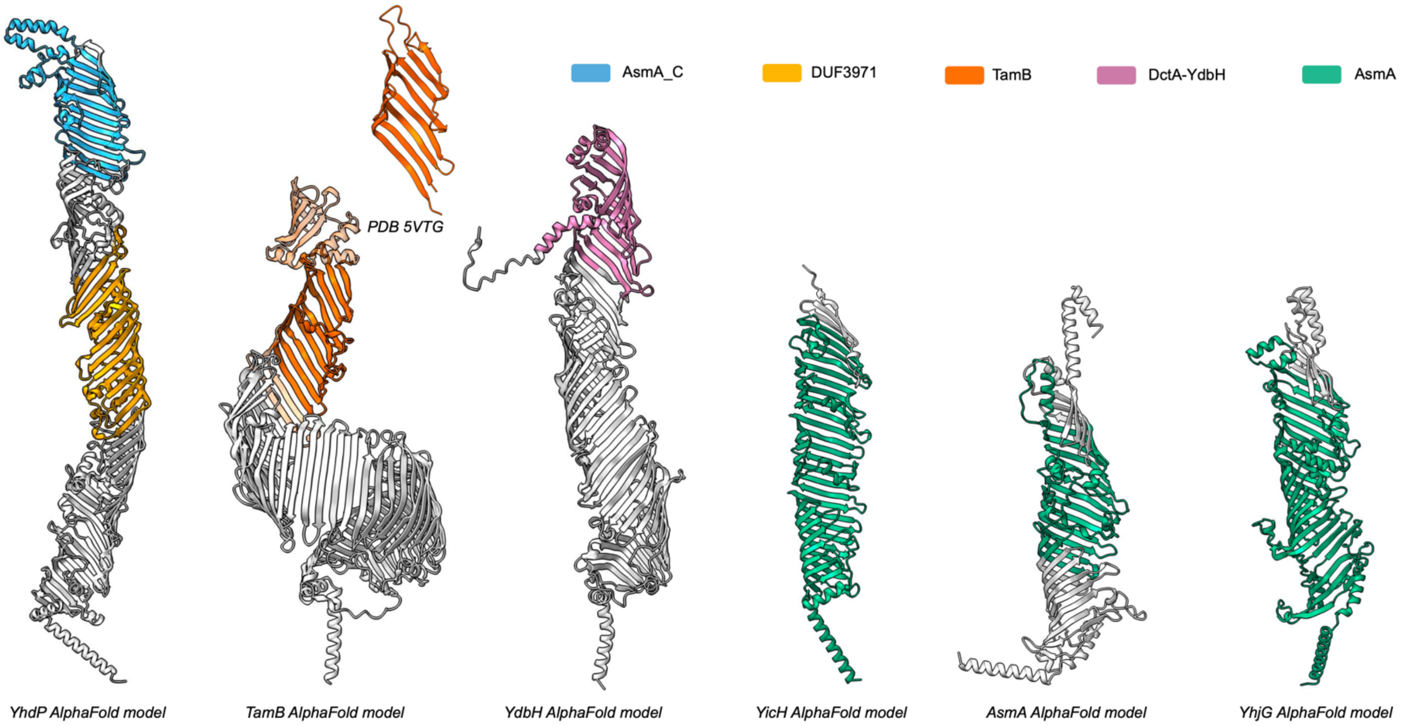


**Supplementary Figure 2 - Annotated domains within the *E. coli* AsmA proteins.** Cartoon representation of the AlphaFold predicted structures for YhdP (AF-P46474-F1), TamB (AF-P39321-F1), YdbH (AF-P52645-F1), YicH (AF-P31433-F1), AsmA (AF-P28249-F1) and YhjG (AF-P37645-F1). Highlighted regions correspond to the domains currently annotated within each of the six proteins. The crystal structure (PDB: 5VTG) of a TamB portion encompassing part of the TamB domain is also shown.

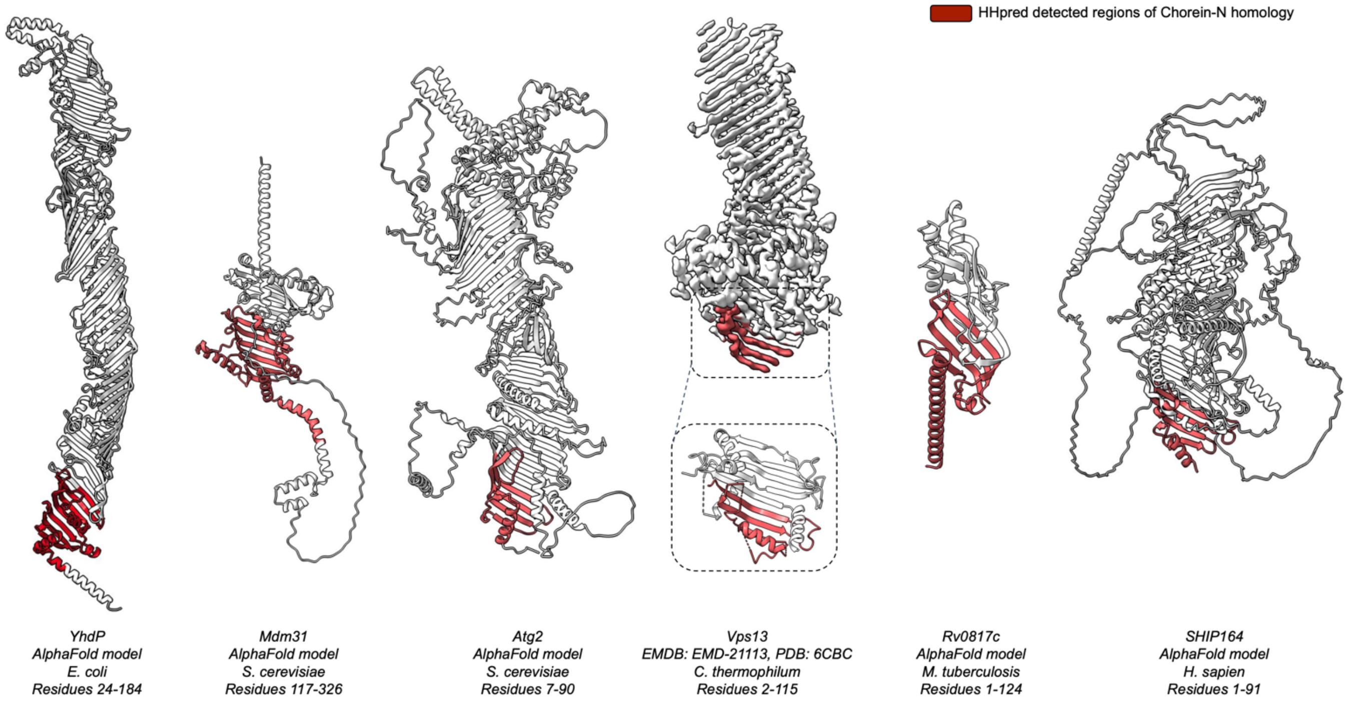


**Supplementary Figure 3 - Sequenced based homology between the *E. coli* AsmA proteins and distant homologs.** Cartoon representation of the AlphaFold predicted structures for YhdP (AF-P46474-F1), Mdm31 (AF-P38880-F1), Atg2 (AF-P53855-F1), Rv0817c (AF-I6WZH9-F1) and SHIP164 (AF-A0JNW5-F1) in addition to a cryo-EM map (EMD-21113) and crystal structure (PDB: 6CBC) of Vps13. Regions of homology, as identified by HHpred, are highlighted in red, whilst residue numbering for the regions are provided beneath each model.

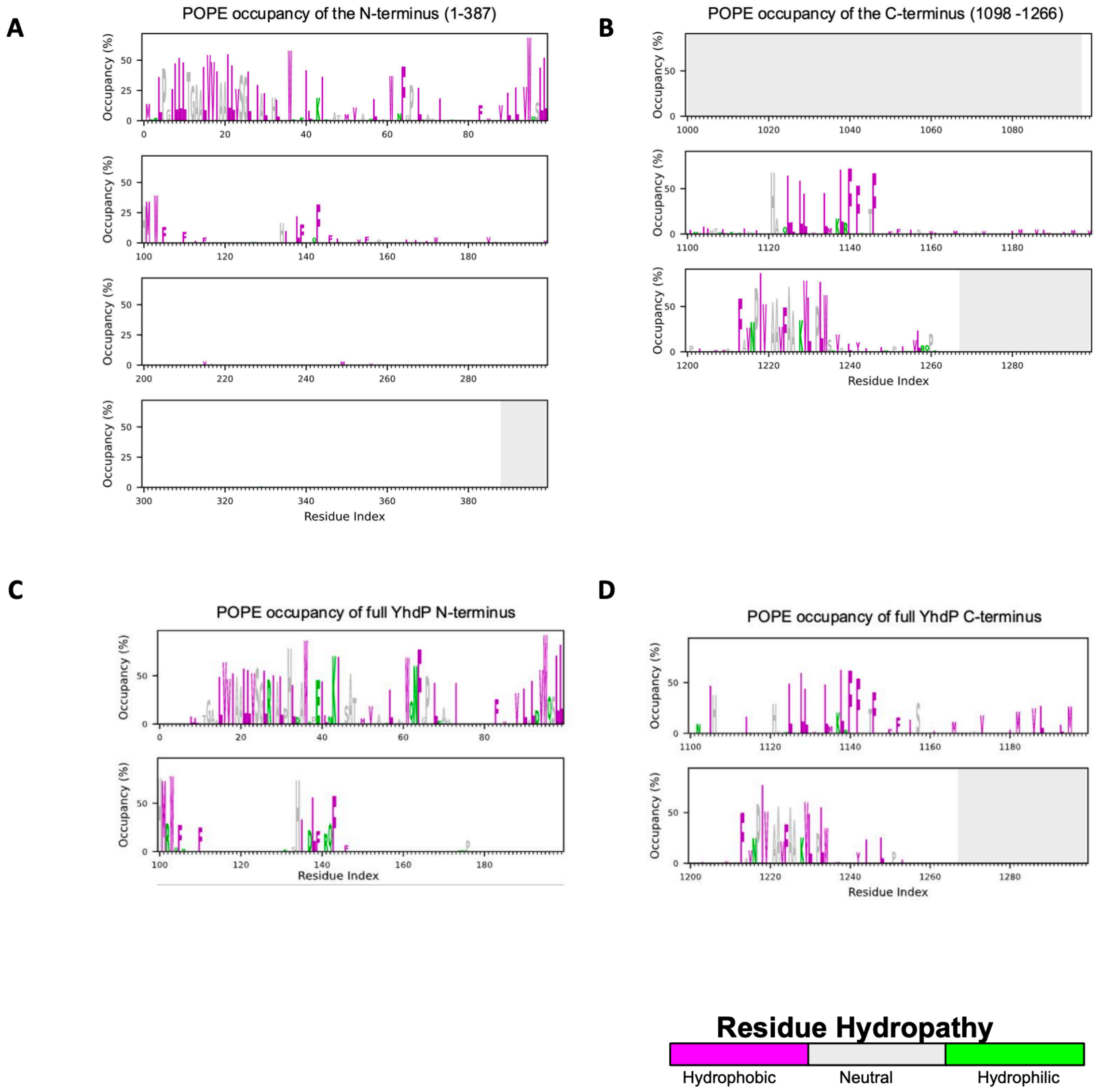


**Supplementary Figure 4 -PyLipID occupancy plots for acyl tail interactions during production simulations.** (A-D) Plots were calculated using PyLipID to measure interactions between YhdP and acyl beads. (A) N-terminal YhdP fragment (residues 1-387) showed acyl tail interactions focused around hydrophobic residues of the TM domain and the line of hydrophobic residues formed by the P-helix and residues W61 and F64. Results are the mean occupancy from 3 simulations, each of 5 μs. (B) C-terminal fragment of YhdP (residues 1098-1266) showed acyl tail interactions that were primarily formed between the hydrophobic residues of Chelix_1 (residues 1202-1238) and Chelix_2 (residues 1120 -1145). Results are the mean occupancy from 5 x 5 μs simulations of the C-terminal fragment. One simulation was not included in the analysis due to highly unlikely placement of the fragment within the membrane (Supplementary Figure 5B). (C-D) Acyl tail interactions within a simulation of full length YhdP in a double membrane. Results are from the mean of 3 x 2 μs simulations. (C) N-terminal acyl tail interactions were mainly localised to hydrophobic residues of the TM domain and the line of hydrophobic residues formed by the P-helix and residues W61 and F64, consistent with the N-terminal fragment simulations. (D) Acyl tail interactions with YhdP at the C-terminus were found primarily with Chelix_1 and Chelix_2, consistent with the C-terminal fragment simulations. Analysis was focused on the most populous lipid species (POPE) within the membrane of interest. Residues are coloured thus: strongly hydrophobic residues are purple, residues which are neutral in terms of hydropathy are grey, and hydrophilic residues are green.

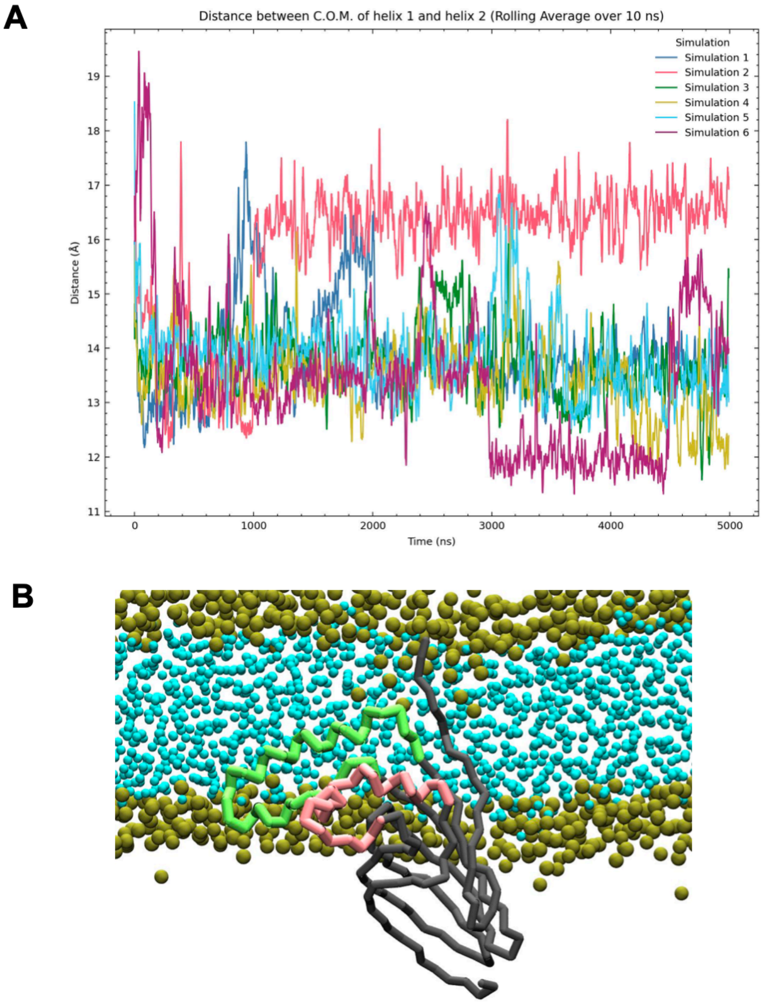


**Supplementary Figure 5 - The C-terminal YhdP fragment (residues 1098-1266) underwent consistent domain movement in 5 of the 6 x 5 μs simulations.** (A) Plot showing the change of distance between centre of masses of 6 simulations of Chelix_1 and Chelix_2 as a function of time. (B) Endpoint snapshot for one of the 6 simulations for the spontaneous assembly of a C-terminal YhdP fragment within a lipid bilayer where the fragment of YhdP was found completely inserted into the membrane, resulting in a different separation of Chelix_1 and Chelix_2 This is common in this membrane assembly procedure due to the stochastic nature of bilayer assembly (86, 87). Backbone shown in dark grey and phosphate beads shown in gold.

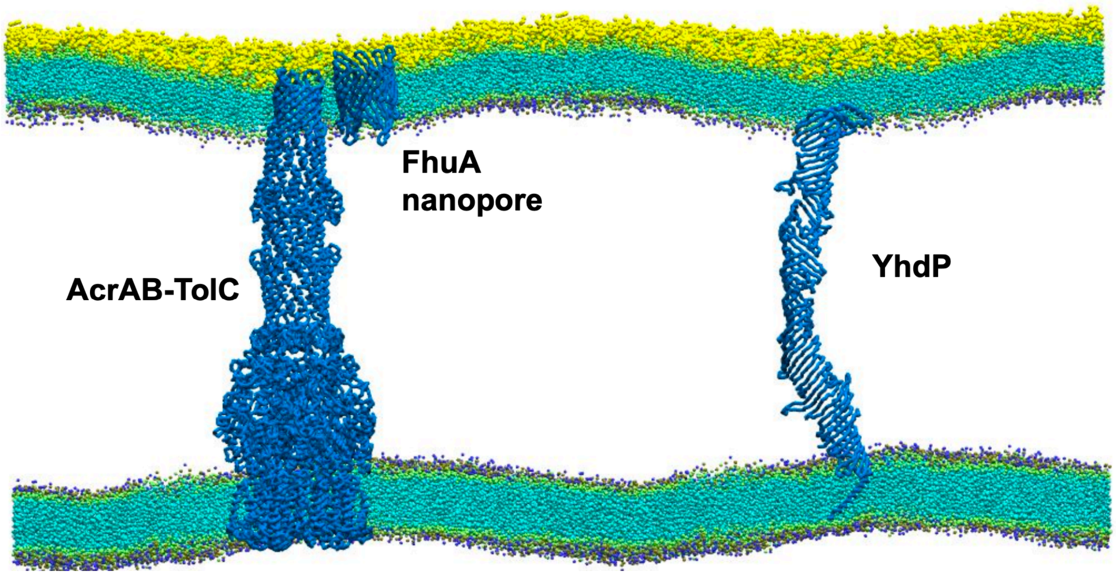


**Supplementary Figure 6 - Snapshot from 3 x 2 μs of YhdP within a double membrane.** The full system consisted of YhdP inserted into our previously assembled cell envelope system comprising AcrAB-TolC, a FhuA nanopore, and a model double membrane (46). Backbone shown in blue. Lipid colour key: acyl beads in cyan, phosphate beads in gold, ethanolamine beads in navy, and sugar beads in yellow.

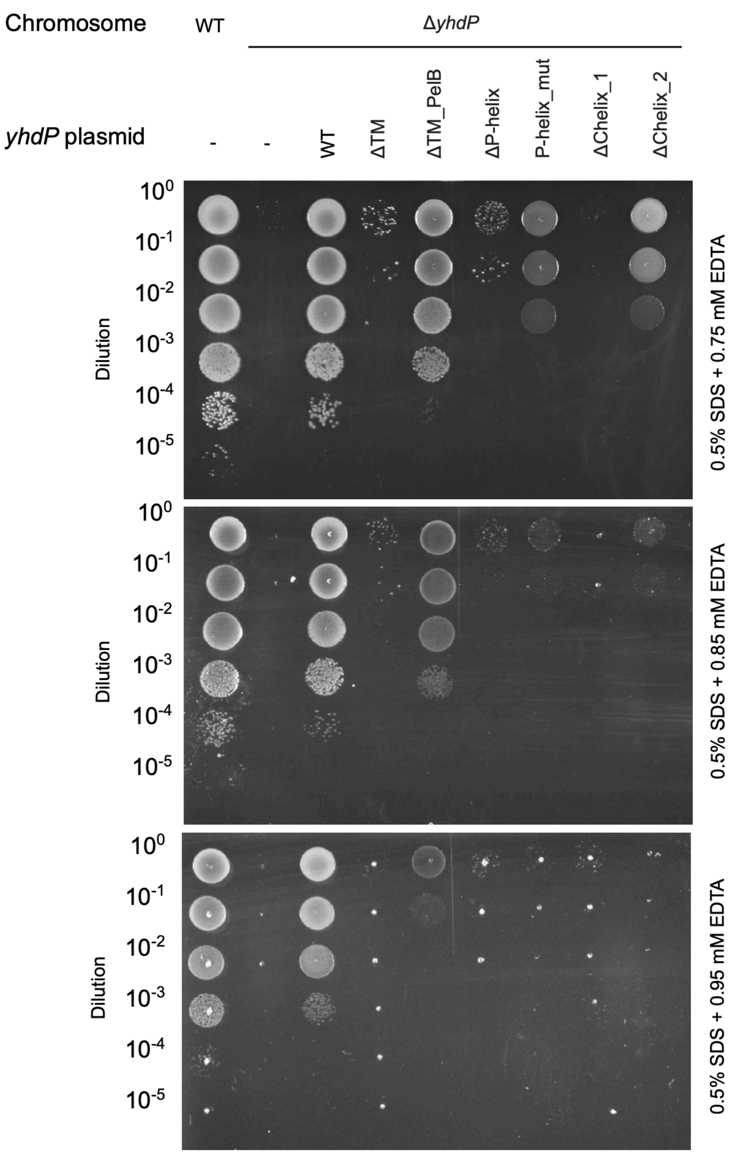


**Supplementary Figure 7 - Genetic complementation of *yhdP* knockout with different mutants of YhdP at varying concentration of EDTA.** 10-fold serial dilutions of the indicated cultures spotted on LB plates containing SDS+EDTA at the concentrations indicated.

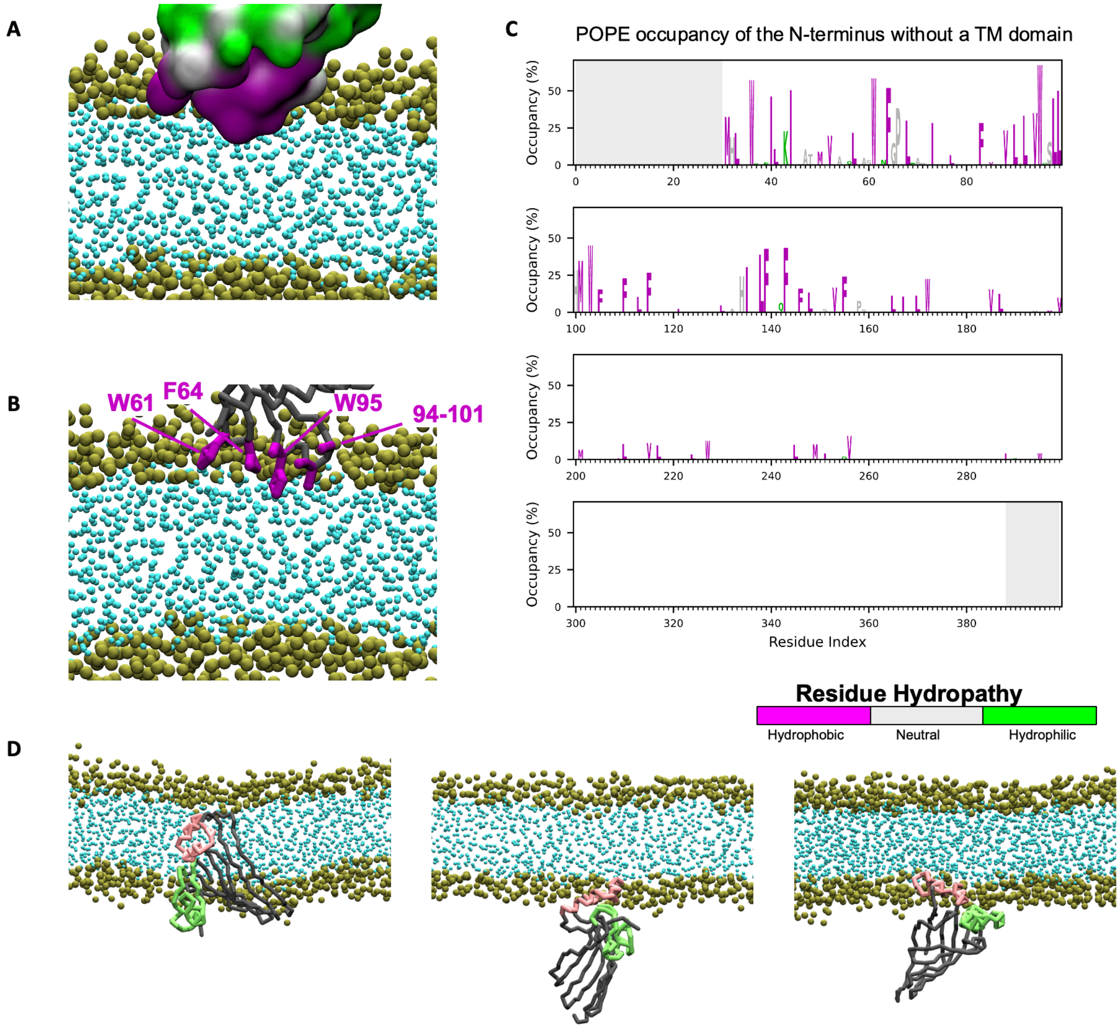


**Supplementary figure 8 - Simulations of mutant fragments of the N- and C-terminus of YhdP**. (A-C) Results from 3 x 5 μs simulations of an N-terminal fragment with TM helix (residues 2-31) removed. Side chains are coloured thus: strongly hydrophobic residues are purple, residues which are neutral in terms of hydropathy are grey, and hydrophilic residues are green. Backbone shown in dark grey. Headgroup beads are shown in gold and acyl tail beads in cyan. (A) Surface representation of the fragment. (B) Consistent with the N-terminal fragment of YhdP with the TM domain, hydrophobic residues W61, F64 and the P-helix residues form a hydrophobic line (shown in sticks) that insert into the membrane, with W95 inserted the most deeply. (C) PylipID occupancy plot of residue interactions with lipid acyl beads. (D) Mutating Chelix_1 residues to SG repeats of the C-terminal fragment (residues 1098-1266) led to random orientation of YhdP during bilayer assembly. YhdP backbone is shown in grey, Chelix_1 (mutated) shown in green, Chelix_2 shown in pink. Headgroup beads are shown in gold and acyl beads in cyan.

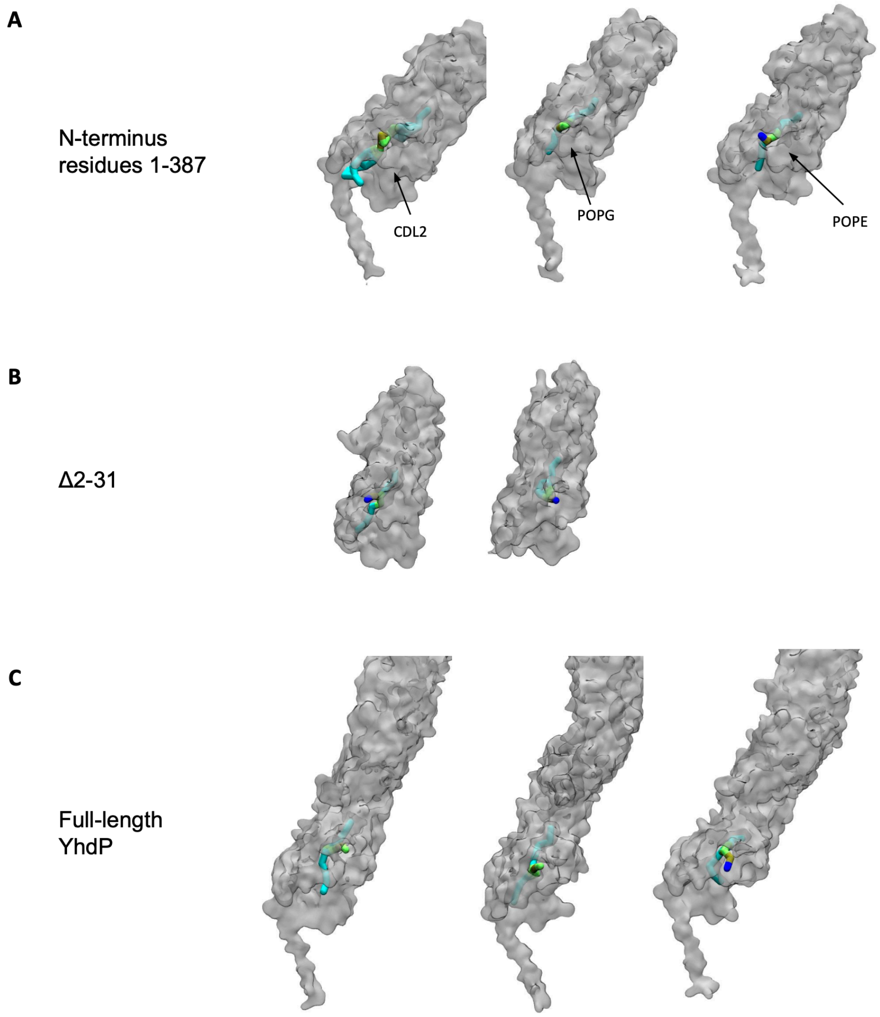


**Supplementary Figure 9 - Lipids entered the hydrophobic groove in 6 of 7 simulations of the N-terminus.** (A-C) Surface representation of a snapshot once lipid had entered the hydrophobic groove. Complete entry was defined as the headgroup emerging from the first cavity. (A) Simulations of the N-terminal YhdP fragment (residues 1-387) had a different lipid enter the hydrophobic groove in each simulation (CDL2, POPG and POPE). (B) POPE moved into the hydrophobic groove during a simulation of the full length YhdP. (C) POPE moved into the N-terminal fragment without the TM domain in 2 of 3 5 μs simulations. Lipid colour key: acyl beads in cyan, ethanolamine beads in navy, glycerol beads in lime, phosphate beads in gold.

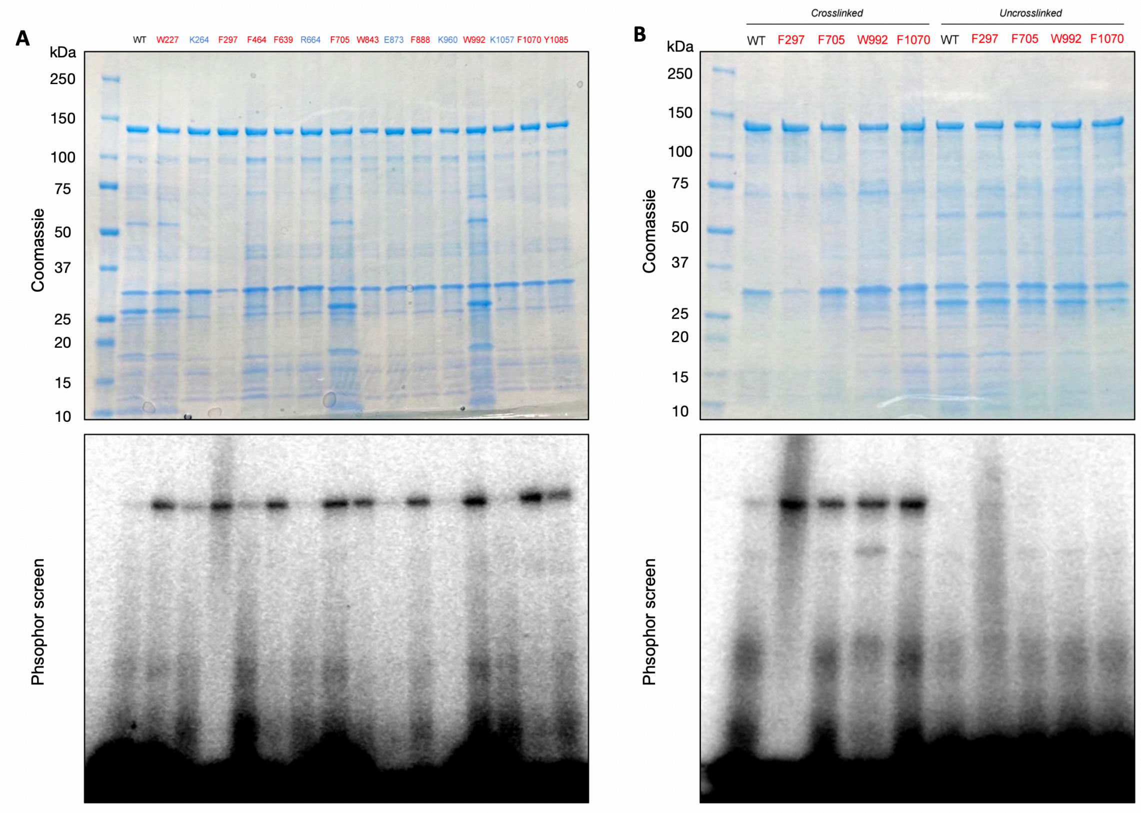


**Supplementary Figure 10 - Uncropped SDS-PAGE gels and phosphor screens of *in vivo* crosslinking assay.** Related to Figure 5.

**Supplementary Video 1 - Full length YhdP remained stably bound to both the inner and outer membrane throughout simulation.** Video from one of the 3 x 2 μs simulations of full length YhdP. A lipid can be seen entering the hydrophobic channel at the N-terminus. Backbone shown in grey, TM domain in cyan, P-helix in yellow, Chelix_1 in green and Chelix_2 in pink. Lipid colour key: acyl beads in cyan, glycerol beads in lime, ethanolamine beads in navy, phosphate beads in gold.

**Supplementary Video 2 - A POPG molecule enters the hydrophobic groove of the YhdP N-terminal fragment.** The phospholipid approaches the YhdP N-terminus and probes the groove. Eventually, the lipid fully enters the groove and the headgroup emerges from cavities in the N-terminal surface. Transparent surface representation of the N-terminal fragment. Lipid colour key: acyl beads in cyan, glycerol beads in lime, phosphate beads in gold. Rest of membrane not shown for clarity.
