## Supplementary tables for "Phospholipid transport to the bacterial outer membrane through an envelope-spanning bridge"

**Supplementary Table 1: plasmids used in this study**

| Plasmid ID | Use of plasmid | Source |
|---|---|---|
| pCP20 | Plasmid for removal of kanamycin resistance cassette | (59) |
| pGLI063 | pET17b derived plasmid containing WT YhdP for complementation | This study |
| pGLI013 | 6xHis-YhdP: pET based plasmid for expression of YhdP with and N-terminal his tag | This study |
| pGLI014 | YhdP-6xHis: pET based plasmid for expression of YhdP with and C-terminal his tag | This study |
| pGLI083 | pET17b-yhdP_phelixmutate, for complementation studies. Hydrophobic residues in the P-helix were mutated: W95T, L98E, L99L, M101N | This study |
| pGLI084 | pET17b-yhdP_phelixdelete, for complementation studies. Deletion of residues 94-103 and replaced with "SG" | This study |
| pGLI085 | pET17b-yhdP_Δctermhelix1, for complementation studies. Deletion of resides 1202-1237 and place with "GSGS" | This study |
| pGLI086 | pET17b-yhdP_Δctermhelix2, for complementation studies. Deletion of resides 1121-1145 and place with "SG" | This study |
| pGLI087 | pET17b-yhdP_Δtmhelix, for complementation studies. Deletion of residues 2-31. | This study |
| pGLI088 | pET17b-yhdP_ΔtmhelixpelB, for complementation studies. Deletion of residues 2-31 and replacement with PelB leader sequence. | This study |
| pGLI067 | pDCE587_yhdp_W227Bpa. Expression plasmid for YhdP W227Bpa mutant for *in vivo* crosslinking | This study |
| pGLI068 | pDCE587_yhdp_K264Bpa. Expression plasmid for YhdP K264Bpa mutant for *in vivo* crosslinking | This study |

| | | |
|---|---|---|
| pGLI069 | pDCE587_yhdp_F297Bpa. Expression plasmid for YhdP F297Bpa mutant for *in vivo* crosslinking | This study |
| pGLI071 | pDCE587_yhdp_F464Bpa. Expression plasmid for YhdP F464Bpa mutant for *in vivo* crosslinking | This study |
| pGLI073 | pDCE587_yhdp_F639Bpa. Expression plasmid for YhdP F639Bpa mutant for *in vivo* crosslinking | This study |
| pGLI074 | pDCE587_yhdp_R664Bpa. Expression plasmid for YhdP R664Bpa mutant for *in vivo* crosslinking | This study |
| pGLI075 | pDCE587_yhdp_F705Bpa. Expression plasmid for YhdP F705Bpa mutant for *in vivo* crosslinking | This study |
| pGLI076 | pDCE587_yhdp_W843Bpa. Expression plasmid for YhdP W843Bpa mutant for *in vivo* crosslinking | This study |
| pGLI078 | pDCE587_yhdp_E873Bpa. Expression plasmid for YhdP E873Bpa mutant for *in vivo* crosslinking | This study |
| pGLI079 | pDCE587_yhdp_F888Bpa. Expression plasmid for YhdP F888Bpa mutant for *in vivo* crosslinking | This study |
| pGLI080 | pDCE587_yhdp_K960Bpa. Expression plasmid for YhdP K960Bpa mutant for *in vivo* crosslinking | This study |
| pGLI081 | pDCE587_yhdp_W992Bpa. Expression plasmid for YhdP W992Bpa mutant for *in vivo* crosslinking | This study |
| pGLI031 | pDCE587_yhdp_F1070Bpa. Expression plasmid for YhdP F1070Bpa mutant for *in vivo* crosslinking | This study |
| pGLI033 | pDCE587_yhdp_Y1085Bpa. Expression plasmid for YhdP Y1085Bpa mutant for *in vivo* crosslinking | This study |
| pGLI034 | pDCE587_yhdp_K1057Bpa. Expression plasmid for YhdP K1057Bpa mutant for *in vivo* crosslinking | This study |
| pEVOL-pBpF | tRNA synthetase/tRNA pair for the in vivo incorporation of a photocrosslinker, p-benzoyl-l-phenylalanine into proteins in E coli in response to the amber codon, TAG. | Chin *et al*, 2002 |

**Supplementary Table 2 - Equilibration and production scheme for N-terminus fragment simulations.**

| STEP | TIME (ns) | TIME STEP (fs) | RESTRAINTS (protein backbone beads; lipid head beads) ( kJ $mol^{-1}$ $nm^{-2}$) | THERMOSTAT | BAROSTAT |
|---|---|---|---|---|---|
| NVT | 1.0 | 2 | 1000; 200 | V-rescale | Berendsen |
| NPT1 | 1.0 | 10 | 250; 50 | V-rescale | Parrinello-Rahman |
| NPT2 | 1.5 | 15 | 100; 20 | V-rescale | Parrinello-Rahman |
| NPT3 | 100 | 20 | 50; 10 | V-rescale | Parrinello-Rahman |
| NPT4 | 40 | 20 | None | V-rescale | Parrinello-Rahman |
| PRODUCTION | 5000 | 20 | None | V-rescale | Parrinello-Rahman |

**Supplementary Table 3 - Equilibration and production scheme for full length YhdP in a double membrane.**

| Step | Time (ns) | Time step (fs) | Thermostat | Barostat |
|---|---|---|---|---|
| NPT1 | 2.5 | 5 | V-rescale | Berendsen |
| NPT2 | 10 | 10 | V-rescale | Parrinello-Rahman |
| NPT3 | 450 | 15 | V-rescale | Parrinello-Rahman |
| PRODUCTION | 2000 | 15 | V-rescale | Parrinello-Rahman |
